# Indexable signal amplification for multiparametric imaging

**DOI:** 10.1101/2021.06.03.446843

**Authors:** Paul D. Simonson, Itzel Valencia, Sanjay S. Patel

## Abstract

Multiparametric imaging allows researchers to measure the expression of many biomarkers simultaneously, allowing detailed characterization of cell microenvironments. One such technique, CODEX, allows fluorescence imaging of >30 proteins in a single tissue section. In the commercial CODEX system, primary antibodies are conjugated to DNA barcodes. This modification can result in antibody dysfunction, and development of a custom antibody panel can be very costly and time consuming as trial and error of modified antibodies proceeds. To address these challenges, we developed novel tyramide-conjugated DNA barcodes that can be used with primary antibodies via peroxidase-conjugated secondary antibodies. This approach results in signal amplification and imaging without the need to conjugate primary antibodies. When combined with commercially available barcode-conjugated primary antibodies, we can very quickly develop working antibody panels. We also present methods to perform antibody staining using a commercially available automated tissue stainer and *in situ* hybridization imaging on a CODEX platform.

## Introduction

Characterization of tumor microenvironments is of great interest for understanding cancer biology and identifying potential targeted therapies^1,2^. While flow cytometry, including CyTOF^3^, allows comprehensive immunoprofiling of cell suspensions, cell-to-cell interactions and other spatial information is lost. Multiparametric tissue imaging, on the other hand, enables characterization of tumor microenvironments with many biomarkers while preserving spatial context^4–6^. Examples of the utility of multiparametric imaging to identify potential targeted disease therapies are multiplying, including CTLA4-blockade for classic Hodgkin lymphoma^7^ and hypomethylating agents for subsets of diffuse large B-cell lymphoma patients^8^.

Current multiparametric imaging platforms vary by maximum number of concurrent targets, throughput, and fundamental methodologies. High throughput platforms like the Opal/Vectra Polaris system (Akoya Biosciences) provide the benefit of signal amplification^5,9^ but are hindered by low multiplexing capacity (generally six or fewer antibodies) and difficulties associated with spectral overlap of the fluorescent molecules used as reporters. More highly multiplexing approaches, including CO-Detection by indEXing (CODEX)^10,11^, Digital Spatial Profiling^12^, cyclic immunofluorescence (CycIF)^13–16^, Imaging Mass Cytometry (Fluidigm)^17^, and multiplexed ion beam imaging by time of flight mass spectrometry (MIBI-TOF)^18,19^, can image more biomarkers but do not allow signal amplification^20^ and are costly, low-throughput, and technically- and labor-intensive. Ultivue InSituPlex is a highly multiplexable solution that allows signal amplification of DNA-barcoded antibodies through the use of a proprietary method^6,21^, though the need for primary antibody conjugation to DNA barcodes remains.

Conjugating antibodies to DNA barcodes, necessary for CODEX and other imaging systems, can result in antibody dysfunction. Development of an antibody panel can therefore be very costly and time consuming as trial and error of modified antibodies proceeds. In some cases, a working barcode-conjugated antibody clone might simply not be identified after great time and expense. For example, in a recent report by Phillips et al.^22^ focused on development of a panel for imaging immunoregulatory proteins by CODEX, a working antibody for CTLA-4 imaging could not be found despite trial of three different clones. Even if a working conjugated antibody is produced, some biomarkers are expressed in such low quantities that the imaging intensity might be insufficient for reliable interpretation. Recently Mistry et al. demonstrated an approach for tyramide signal amplification in CODEX^23^ with three amplified marker signals (presumably one for each of the available microscope fluorescence channels), and the imaging was performed after completion of the normal CODEX imaging. This approach, while useful, is limited due to the inability to go beyond the number of available fluorescence channels and the need for removing the sample from the imaging system for additional staining and re-imaging.

In this report, we present a generalizable method for CODEX imaging with unmodified primary antibodies and signal amplification that uses novel tyramide-conjugated DNA barcodes (tyramide-barcodes). This allows us to use the benefits of tyramide signal amplification, but image the targets in an indexed way using a commercial CODEX imaging system (now rebranded as PhenoCycler™, Akoya Biosciences). This approach is flexible, and we also demonstrate that it can be used with *in situ* hybridization probes. In order to make this system workable, we also developed a staining approach that allows us to perform both tyramide-barcode staining and standard CODEX staining using a commercially available automated tissue stainer, made possible using a custom coverslip holder. This drastically reduces the amount of manual labor required for CODEX staining and has the potential to improve staining reproducibility.

## Results

### Staining automation

In order to integrate tyramide-barcode staining with regular CODEX staining using barcode-conjugated primary antibodies, we determined it was desirable to automate the staining of tissue samples for CODEX imaging. In the commercial version of CODEX, antibody staining is carried out manually using tissue attached to a coverslip. A coverslip is required for the optical imaging, yet commercial automated stainers are generally designed to work with microscope slides. We therefore developed a custom metal coverslip adapter (Fig. S1), which enabled us to stain coverslips on a commercial tissue autostainer. We were then able to adapt the manual CODEX staining protocols to the autostainer, with final fixation of antibodies to the tissue performed manually after removal of the coverslip from the autostainer.

### Tyramide-barcode staining is compatible with CODEX imaging

In the tyramide-barcode staining approach, primary antibody is incubated with tissue, followed by secondary (or secondary followed by tertiary) antibody that is conjugated to peroxidase. The sample is then incubated with tyramide-conjugated barcodes (tyramide-barcodes). These steps are performed in the autostainer and repeated for additional primary antibodies, after which standard CODEX staining with a cocktail of barcode-conjugated primary antibodies can also be performed (See Fig. 1 and Methods). Acceptable staining is defined as achieving staining that is specific to the intended target (for example, PAX-5 staining demonstrates specific staining in nuclei of B cells) without unacceptably increased background signal or off-target staining.

**Figure 1.**
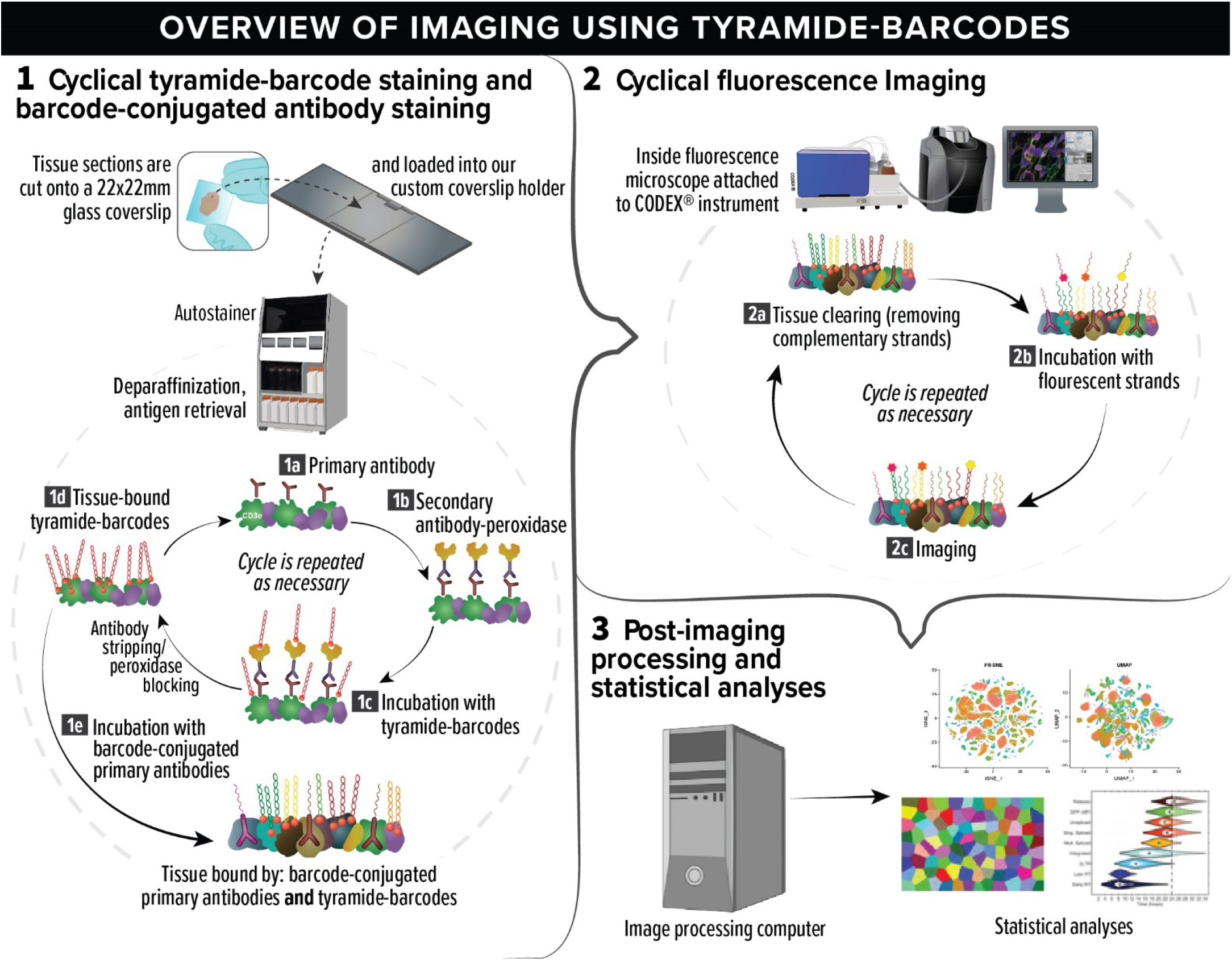
Multiparametric imaging with tyramide-barcodes and barcode-conjugated antibodies. The process can be divided into three stages: (1) tissue staining with tyramide-barcodes and barcode-conjugated antibodies, (2) cyclical fluorescence imaging, and (3) image processing and statistical analysis. In stage 1, tissue is cut into thin sections and placed on microscope coverslips. The coverslips are then loaded onto our custom coverslip holders, which allow processing in a tissue autostainer. Paraffin-embedded tissue is deparaffinized, after which antigen retrieval is performed. Cyclical tyramide-barcode staining is then repeated (steps 1a to 1d) for as many primary antibodies as desired for signal amplification and/or unmodified primary antibody use. In step 1a, the tissue is incubated with an unmodified primary antibody. In step 1b, the tissue is incubated with peroxidase-conjugated secondary antibodies. In step 1c, the peroxidase (attached to the secondary antibodies) enzymatically activates the tyramide, and tyramide-barcodes bind to the tissue in close proximity to the peroxidase. The peroxidase is then permanently inactivated and/or the antibodies are removed, and the cycle can begin again at step 1a for a different primary antibody, or, if there are no more primary antibodies for signal amplification, staining proceeds to step 1e. In step 1e, the tissue is incubated with a cocktail of many barcode-conjugated primary antibodies. The tissue is lightly fixed to lock antibodies into place, and the process then moves on to stage 2. In step 2a, “tissue clearing” is performed, in which double-stranded DNA barcodes are dissociated, and the non-covalently attached strands are washed away. In 2b, one to three unique single-stranded DNA barcodes are incubated with the tissue, each one conjugated to one of three different fluorophores and complementary to one of the tissue-attached barcodes. After incubation, the tissue is imaged by fluorescence imaging, step2c. By imaging only a small subset of the barcodes in each cycle, high quality imaging of each target antigen can be achieved without significant spectral overlap of fluorescence signals. Steps 2a-2c are repeated until all of the antigens have been imaged. Stage 3 then commences, comprising post-imaging image alignment, stitching, and segmentation, followed additional study-specific statistical analyses.

As proof of principle, we began by imaging one, then two, antigens within a single tissue section (for example, see Fig. S2). To evaluate for possible non-specific staining as a result of the introduced barcodes, comparison with tissue stained with tyramide conjugated directly to Cy3 and Cy5 fluorophores instead of barcodes was performed, which demonstrated no noticeable difference in localization or specificity of staining (see Fig. S5). We then demonstrated that we could cycle tyramide-barcode fluorescence on and off, with appropriate clearing between imaging cycles, as is required for CODEX imaging. We did this by imaging a single antigen, CD3e in tonsil tissue, stained using a single tyramide-barcode. We imaged the CD3e in two cycles on a CODEX system. In the first cycle, we imaged using complementary strands attached to Cy5 fluorophore. In the next cycle, we imaged using complementary strands attached to Cy3. We saw appropriate clearing and staining for both cycles (see Fig. S10). Importantly, we found it necessary to use non-fluorescent tyramide-conjugated DNA barcodes when incubating with peroxidase in the autostainer; otherwise, bright fluorescence would be present throughout imaging, despite tissue clearing. This finding suggested that in fact a significant amount of tyramide-conjugated double-stranded DNA barcode was attached to the tissue in such a way that some of the DNA could not be dissociated and washed away, under the tested conditions.

In order to directly demonstrate amplification of signal by tyramide barcodes in comparison to standard CODEX imaging, we stained tonsil tissue using preconjugated CD3e antibody (clone EP449e, Akoya Biosciences, barcode 45 with Cy5 reporter) and performed standard CODEX imaging with the complementary barcode. We also stained a section of the same tonsil using the same primary antibody but included steps for amplification using tyramide-barcode; imaging was performed using Cy5-conjugated reporter that was complementary to the tyramide barcode. Staining localization was localized to T cells and appropriate for both stained tissue sections, but fluorescence signal intensity was approximately 4.7x greater for the tyramide-barcode stained tissue (see Fig. S6). Image resolution also appeared similar for the two experiments (Fig. S6).

Given these successes, we extended the tyramide-barcode staining technique to imaging using *in situ* hybridization probes for detection of RNA. Epstein-Barr-virus-encoded small RNAs (EBERs) are noncoding RNAs that are expressed abundantly in EBV-infected cells, and *in situ* hybridization probes are commonly used to detect them in clinical laboratories. The *in situ* hybridization probes are often conjugated to FITC at one end, and anti-FITC primary antibodies, followed by secondary, then tertiary peroxidase-conjugated antibodies, are used for detection using a chromogenic substrate (catalyzed by peroxidase). We performed tyramide-barcode staining by simply replacing the chromogenic substrate with tyramide-barcode, and we tested the system using a sample of classic Hodgkin lymphoma tissue wherein the Hodgkin cells were known to be infected with EBV. Our system clearly highlighted Hodgkin cells (Fig. S11) and was appropriately negative for negative controls. These findings demonstrate the flexibility of the tyramide-barcode approach, beyond just protein detection via primary antibodies, for CODEX imaging.

We then demonstrated tyramide-barcode staining with multiple primary antibodies. This required multiple staining cycles, one for each unique primary antibody, with inactivation of peroxidase prior to incubation with each subsequent primary antibody. Staining lymph node tissue with four primary antibodies (CD3e, CD4, CD10, and ICOS) in this manner demonstrated appropriately localized staining (see Fig. S12).

### Integration with standard CODEX antibodies

We next incorporated tyramide-barcode staining with multiple primary antibodies into a complete CODEX workflow so that we could stain some markers with tyramide-barcodes and some with commercially available barcode-conjugated primary antibodies. All deparaffinization, antigen retrieval, and staining steps were performed in a Leica Bond RX autostainer using tissue attached to a coverslip and our custom coverslip holder. CD3e and CD20 antigens were stained using commercially available DNA barcode conjugated primary antibodies, and CD4 and CD163 antigens were stained using our tyramide-barcodes staining approach. Example images demonstrating the results of the integrated staining protocol are seen in Fig. 2, which demonstrates appropriately localized staining for each of the target antigens. Since the peroxidase should be removed (e.g., by heat-induced stripping away of tissue-bound antibodies that are conjugated to peroxidase) or inactivated (e.g., by application of a solution containing hydrogen peroxide) before staining with each new tyramide barcode, we attempted combinations of treatment with hydrogen peroxide solution and heat-induced antibody stripping. For this particular experiment, we found that simple peroxidase inactivation with hydrogen peroxide solution produced better signal intensity with adequately reduced off-target staining. The better signal intensity we saw is hypothesized to be due to better preservation of the tissue integrity (since heat-induced antibody stripping was not applied) and/or better maintenance of attachment of tyramide bound to tissue in earlier steps; such protocol optimizations are also commonly necessary for other methods that use multiple rounds of tyramide staining, such as that used in staining for imaging on a Vectra Polaris instrument^5^. The final protocol for this particular experiment, which is that used to produce the images seen in Fig. 2, is found under “Combined tyramide-barcode and barcode-conjugated primary antibody staining” in the Methods section.

**Figure 2.**
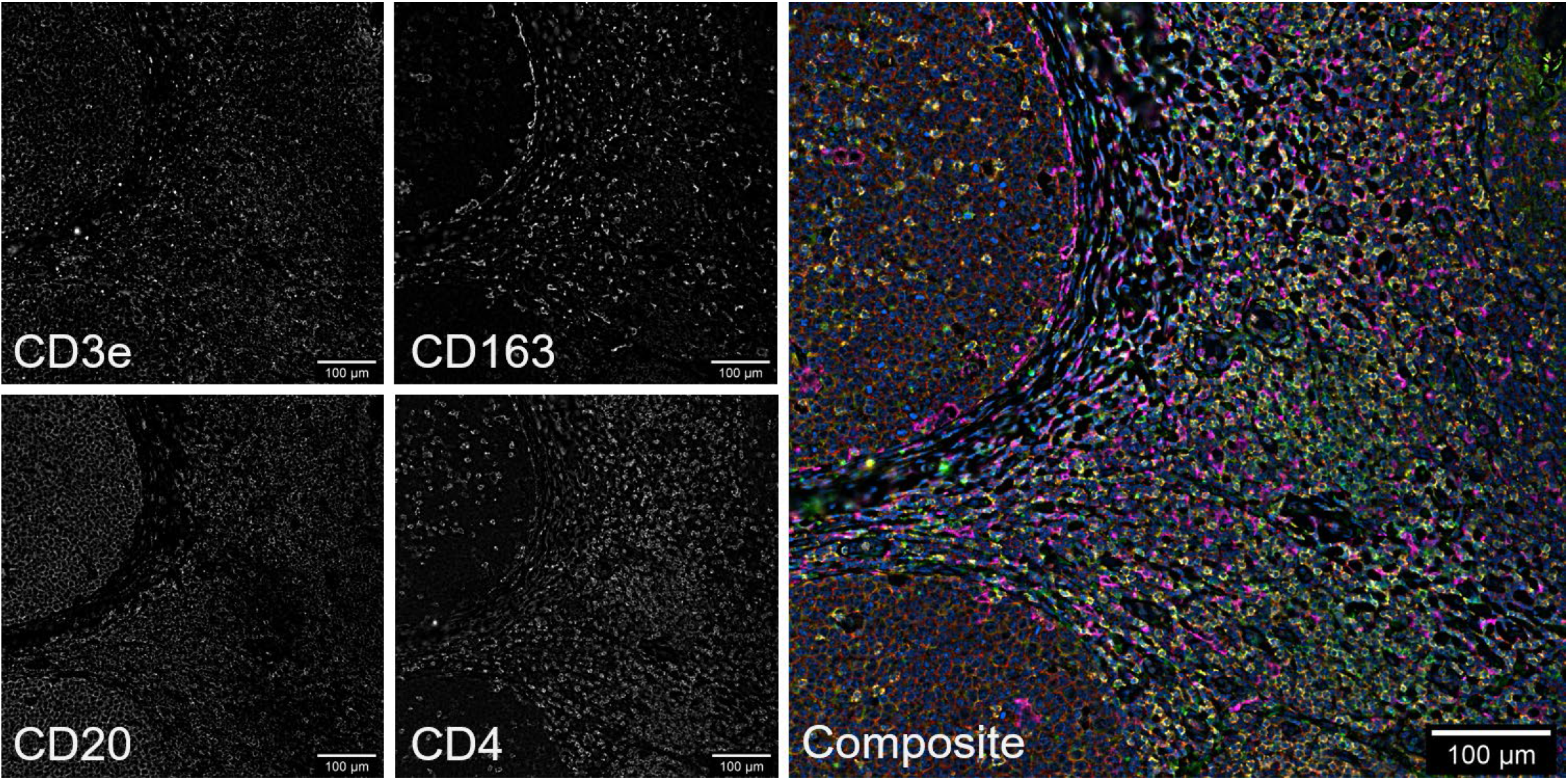
Combined tyramide-barcode and conjugated primary antibody staining in tonsil. CD3e (1:100 dilution) and CD20 (1:200 dilution) were stained using commercial barcode-conjugated primary antibodies (Akoya Biosciences), and staining of CD4 and CD163 was performed using tyramide-barcodes with mouse anti-CD4 (1:100 dilution, clone 4B12, Leica Biosystems) and mouse anti-CD163 (ready-to-use format, clone 10D6, Leica Biosystems) primary antibodies. The composite image is colored as follows: CD20 = red, CD4 = yellow, CD3e = green, and CD163 = magenta. The Keyence BZ-X800 microscope used 100% excitation intensity, “high resolution” imaging mode, and a Nikon 20x PlanApo lambda 0.75 NA objective. Background subtraction and image stitching were performed using Akoya’s CODEX Processor software.

### Rapid antibody panel development for classic Hodgkin lymphoma

As an additional example of the use of the full method, we imaged classic Hodgkin lymphoma using a custom CODEX antibody panel that was comprised of seven commercially available barcode-conjugated primary antibodies, and also included antibodies for two additional markers, CD30 and MUM1, for which commercial CODEX barcode-conjugated antibodies were not available. MUM1 is a protein that is very frequently positive in the nuclei of Hodgkin/Reed-Sternberg cells. CD30 staining is routinely used in diagnostic clinical practice to identify Hodgkin/Reed-Sternberg cells, and thus comparison with tissue that is already stained for CD30 expression by immunohistochemistry is readily available. Furthermore, as it is quite specific for the Hodgkin cell membranes with respect to the surrounding cell milieu, it has the potential to be very useful for simplifying cell image segmentation (though we do not report this approach here). Prior experience at the time in attempting to create a barcode-conjugated CD30 antibody demonstrated little to no staining after conjugation, despite trying two antibody clones (Ber-H2, BioLegend, cat. no. 333902; Ber-H8, BioRad, cat. no. MCA6134). Therefore, we decided CD30 would be a particularly good target for using tyramide-barcode staining for imaging by CODEX.

Staining was performed as detailed in the Methods section. As a notable difference in the staining protocol of this experiment compared to that of the prior section, we opted to incubate the sample with preconjugated antibodies overnight at 4°C and found specific staining with less volume of antibody solution required (the former protocol required two 90 minute incubations with antibodies to avoid the sample becoming dried out in the autostainer, which increased the volume of antibody needed; autostainers also have a “dead volume” which requires the use of more conjugated antibody solution, which is an expensive component of these experiments). By staining CD30 and MUM1 antigens using tyramide-barcodes, we were able to image all the antigens on a CODEX system with no custom antibody conjugation and thus very little panel development effort. DAPI and eosin fluorescence imaging was performed using the CODEX instrument at the end of the imaging run to perform virtual H&E imaging^24^ for morphologic comparison with CODEX images in the same tissue section. The staining thus achieved demonstrated that the imaged markers were appropriately localized, with strong staining of CD30 and MUM1 seen in the cell membranes and nuclei, respectively, of Hodgkin/Reed-Sternberg cells (see Fig. 3 and additional images in Fig. S13-S18). The localization of the fluorescence staining also correlated well with the morphology seen by virtual H&E staining, as assessed by two board-certified hematopathologists. While not the focus of this paper, we also performed image segmentation and spatial analysis (Figs. S15-S18) to review the localization and interaction of various cell types using CODEX analysis software (Akoya Biosciences). This analysis highlighted the strong staining seen by the tyramide-barcodes approach and compatibility with existing software.

**Figure 3.**
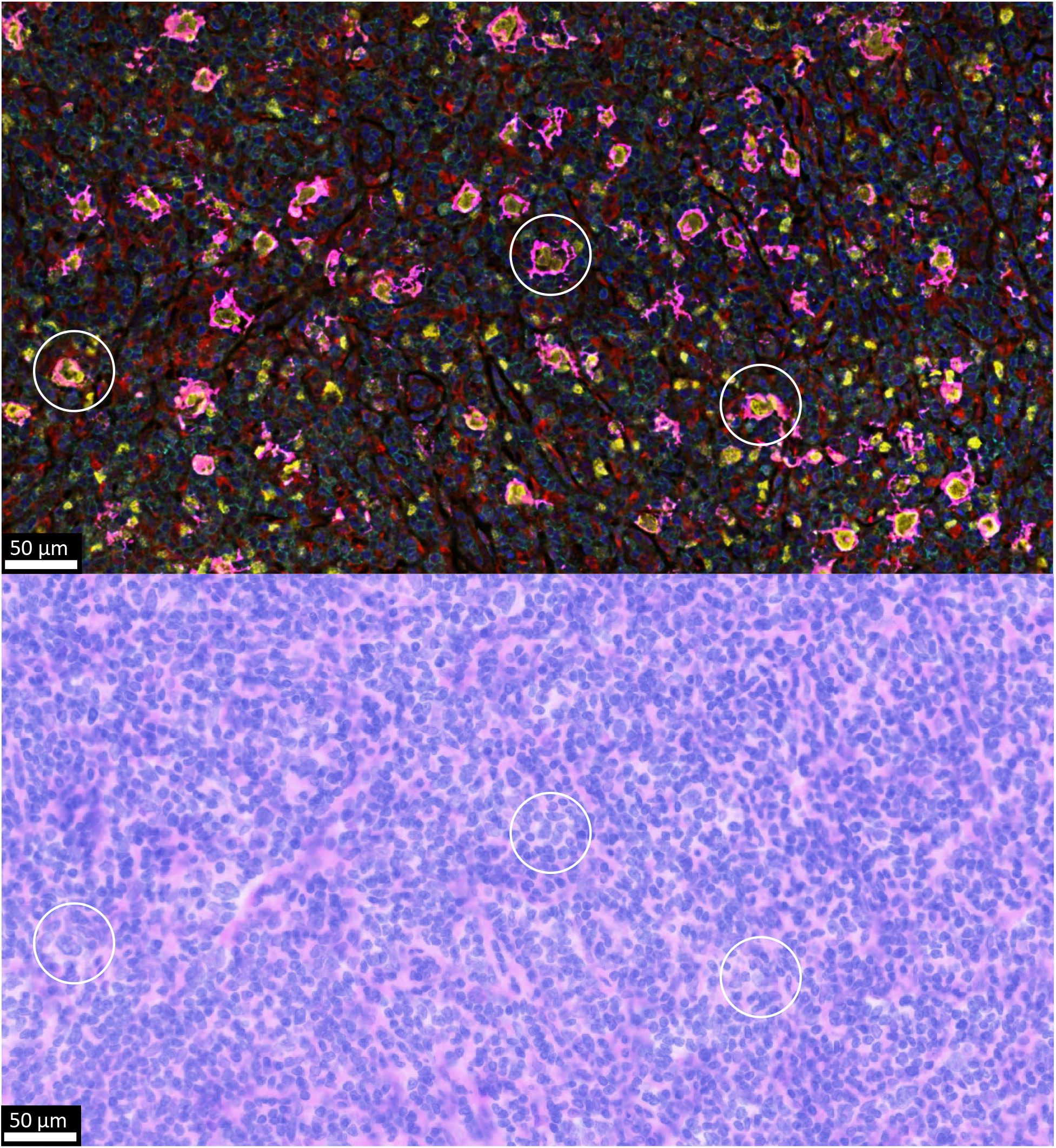
Classic Hodgkin lymphoma stained using a combination of tyramide-barcodes and barcode-conjugated primary antibodies. CD30 (1:50 dilution, mouse host, clone Ber-H2, Invitrogen) and MUM1 (1:100 dilution, mouse host, clone MUM1p, Agilent Dako Products) staining was performed using the tyramide-barcodes, which stains the cell membranes and nuclei of Hodgkin cells, respectively (scattered T cells also express MUM1). Other antigens were stained using commercially available barcode-conjugated primary antibodies (Akoya Biosciences, 1:200 dilutions except for CD3e, which was 1:100 dilution). DAPI = blue, CD30 = magenta, MUM1 = yellow, CD68 = red, CD3e = cyan. Not shown: CD4, CD8, CD11c, CD20, CD45RO, CD107a, and Ki67. Virtual H&E staining (bottom) was generated using DAPI and eosin fluorescence images as described previously^24^. Circles help to identify Hodgkin/Reed-Sternberg cells seen in the fluorescence image and corresponding virtual H&E image.

## Discussion

Standard CODEX imaging makes use of primary antibodies conjugated with DNA oligomers (barcodes), which are in turn hybridized with complementary fluorophore-conjugated DNA strands^11^. Fluorescence imaging is performed in cycles, with only a few complementary strands imaged per cycle so as to avoid the problem of spectral overlap that would be had if many fluorophores were used^10,11^. Cycling continues until all of the antigen types, indexed by barcodes, have been imaged (see stage 2 in Fig. 1). In the normal situation, there is no signal amplification since the number of barcodes is proportional to the number of bound primary antibodies. Thus, this approach is limited.

In this work, we have developed new reagents, tyramide-barcodes, that can be used to stain tissue using unmodified primary antibodies and achieve signal amplification. Tyramide-barcodes are flexible and can in theory be used in any peroxidase-based signal amplification system, including in *in situ* hybridization imaging approaches (see Fig. S4). Use of barcodes means that in theory a large number of signals can be amplified, including when using primary antibodies from the same host species, assuming appropriate blocking/removal of peroxidase can be achieved between staining cycles similar to that required for such systems as the Vectra Polaris.

Besides signal amplification, perhaps the greatest benefit of this new approach is the lack of need for primary antibody conjugation with DNA barcodes. This approach is compatible with any primary antibody, including those supplied in solutions with carrier protein (e.g. bovine serum albumin), which normally must be absent when conjugating antibodies to DNA barcodes or lanthanide metals (for Imaging Mass Cytometry). Thus, it is possible to use virtually all commercially available antibodies used for immunohistochemistry, with no need for additional purification. Moreover, primary antibodies previously validated by a given lab should be compatible, obviating the need to purchase new reagents and enabling the lab members to capitalize on their prior expertise with a given antibody clone. That experience can help direct optimization of the staining protocol, which is in fact significant given that the staining conditions (time, concentration, etc.) can be customized for each antibody. In our own experience, development of a custom CODEX antibody panel can be significantly hindered by just a few troublesome biomarkers for which a working barcode-conjugated antibody clone is difficult to find. By instead using the tyramide-barcode staining approach, as we did for CD30 staining in classic Hodgkin lymphoma, experiments can go forward without an inordinate amount of time and money spent on the few troublesome biomarkers. The net result is increased efficiency at a reduced cost.

There are limitations to the tyramide-barcode staining approach. One potential limitation could be steric blocking by tyramide-barcodes of antibodies to antigens in later staining steps. For example, in T cells, if CD3e staining signal is amplified in an early staining step, it is possible that CD4 staining in a later step would be reduced due to the CD3-associated barcodes blocking the binding of the CD4 antibodies. We note, however, that in Vectra Polaris imaging, this potential steric hindrance is not generally observed to be an obstacle. Another potential limitation is that the ability to reliably quantify the amount of antigen present is reduced since the amplified signal is not necessarily linearly proportional to the amount of bound primary antibody (similar limitations exist for other tissue signal amplification staining methods, such as immunohistochemistry). The approach is also limited by the amount of time the staining takes to complete since for each additional primary antibody used for amplification, the process can take an additional 1-2 hours to complete (hence, we normally perform the automated tissue staining overnight). Excessive peroxidase blocking and/or removal of antibodies between tyramide-barcode amplification cycles could also lead to increased tissue deterioration for downstream biomarker staining. Hence, limiting the number of amplified biomarker signals is still considered advantageous in that it saves staining time and reduces possible tissue deterioration. Finally, the achievable image resolution is expected to be reduced due to tyramide binding at sites distant from the primary antibody. The decrease in resolution is unlikely to be noticeable in most imaging applications (and was not noticeable in our experiments; see also Fig. S6) but should be noticeable in super-resolution imaging.

In summary, we have developed tyramide-conjugated DNA barcodes that can be used with unmodified primary antibodies in a CODEX imaging system. These enable amplification of the staining signal and imaging of biomarkers for which barcode-conjugated primary antibodies are difficult to generate. We have developed a custom coverslip holder and associated automated CODEX staining workflow, which also allows combining both tyramide-conjugated barcodes and standard barcode-conjugated primary antibodies in a single workflow. Future work will include further expansion and refinement of the staining protocols, use of additional fluorophores, and further incorporation of *in situ* hybridization probes. We will apply the approach to many more imaging applications as we seek to further understand cellular microenvironments and their potential utility for predicting clinical outcomes and response to therapy.

## Methods

Additional methods can be found in the Supplementary Information.

### Construction of tyramide-barcodes

DNA barcode sequences were selected from a list of oligomers that were previously reported as successfully used in CODEX imaging^11^. For one construct, we created the following custom-conjugated oligomer (5’ to 3’, custom orders to Gene Link, Elmsford, New York):

/tyramide/C10/CAAGGAACTACCGA

where C10 is a linker consisting of a chain of 10 carbon atoms. Tyramide was attached to the linker via an NHS ester linkage. Complementary DNA oligomers were also made, with and without attached fluorophores at the 5’ end. Stock solutions were made by diluting oligomer to 1 mg/ml in TE pH 8.0 buffer and stored at 4°C or less.

### Autostainer coverslip holders

The coverslip holder was designed using FreeCAD software (see Fig. S1) and manufactured using CNC machining (ProtoLabs, Maple Plain, Minnesota). The long edges were outlined by a hydrophobic pen (Hydrophobic Barrier PAP Pen, Vector Laboratories) to reduce the chances of leakage in the instrument, though significant leakage has not yet been observed. Before placing a coverslip on the coverslip holder, a small droplet of PBS buffer is placed to help the coverslip adhere. The coverslip holder is washed with water and ethanol between uses.

### Combined tyramide-barcode and barcode-conjugated primary antibody staining

For the imaging seen in Fig. 2, de-identified tonsil tissue was cut into 4-micron-thick sections and placed on charged coverslips. Using a Leica Bond RX automated stainer and our custom coverslip holder, the tissue was deparaffinized, followed by antigen retrieval (Leica ER2 solution, 20 minutes). The protocol then proceeded as follows with Bond Wash Solution (Leica) used for washing: Leica peroxidase blocking solution (5 minutes), wash 3x, mouse anti-CD4 primary antibody (clone 4B12, Leica Biosystems, catalog number NCL-L-CD4-368) diluted 1:100 in Abcam antibody diluent (30 minutes), wash 3x for 2 minutes each, Leica rabbit anti-mouse antibody (8 minutes), wash 3x for 2 minutes each, Leica anti-rabbit antibody conjugated with HRP (8 minutes), wash 3x for 2 minutes each, double-stranded tyramide-barcode #76^11^ (30 minutes), wash (5 minutes), Leica peroxidase blocking solution (5 minutes), wash 3x, mouse anti-CD163 (clone 10D6, Leica Biosystems, catalog number PA0090) ready-to-use solution (30 minutes), wash 3x for 2 minutes each, Leica rabbit anti-mouse antibody (8 minutes), wash 3x for 2 minutes each, Leica anti-rabbit antibody conjugated with HRP (8 minutes), wash 3x for 2 minutes each, double-stranded tyramide-barcode #79^11^ (30 minutes), de-ionized water 2x for 10 minutes each, Akoya hydration buffer 2x for 2 minutes each, Akoya staining buffer (25 minutes), conjugated primary antibodies cocktail (Akoya CD3e 1:100 and Akoya CD20 1:200 in Abcam Antibody Diluent) 2x for 90 minutes each, wash.

The coverslip was then washed 2x with Akoya Staining Buffer for 2 minutes each before being placed in 1.6% formalin in Akoya Staining Buffer for 10 minutes. The coverslip was then washed 3x in PBS before being placed in cold methanol for 5 minutes. The coverslip was then washed 3x in PBS before being placed briefly in Akoya Storage Buffer. Imaging then proceeded as usual for CODEX imaging using the protocols outlined in the Akoya CODEX manual (revision C), but with solutions of 400 nM Cy5-conjugated complementary barcode used as appropriate for the tyramide-barcode stained markers (CD4 and CD163). At the conclusion of CODEX imaging, we used a weak solution of DAPI and eosin to acquire fluorescence images to generate virtual H&E images^24^.

### Antibody staining of classic Hodgkin lymphoma

Staining of classic Hodgkin lymphoma (Fig. 3) proceeded as outlined in the section above, with a tyramide-barcode staining cycle performed for mouse anti-CD30 primary antibody (1:50 dilution, clone Ber-H2, Invitrogen, catalog number MA5-13219), and a tyramide-barcode staining cycle performed for mouse anti-MUM1 primary antibody (1:100 dilution, clone MUM1p, Agilent Dako Products, catalog number GA64461-2). However, in this case the tissue was incubated with the barcode-conjugated primary antibodies cocktail in a hydration chamber at 4°C overnight rather than inside the autostainer at room temperature for two 90 minute cycles. The barcode-conjugated primary antibodies included CD68, CD3e, CD4, CD8, CD11c, CD20, CD45RO, CD107a, and Ki67 (Akoya Biosciences, 1:200 dilution each, except CD3e which was 1:100 dilution).

### Imaging

A commercial Akoya CODEX instrument and a Keyence BZ-X800 microscope with a 20x Nikon PlanApo NA 0.75 objective were used to clear tissue and stain with complementary fluorescent barcodes. The standard Akoya CODEX protocols were followed (Akoya CODEX manual, revision C). For tyramide-barcode staining, we replaced the Akoya barcodes in the reporter plate solutions with our custom complementary barcodes, with a final concentration of 400 nM for each. Post-acquisition image processing was performed using the Akoya CODEX Processor software, and inspection of results was performed using ImageJ with the Akoya CODEX MAV plugin.

## Acknowledgements

This work was supported by the Department of Clinical Pathology and Laboratory Medicine at Weill Cornell Medicine. We thank Giorgio Inghirami and Massimo Loda for review of the manuscript and constructive comments prior to submission for publication.

## Contributions

P.D.S. developed the tyramide-oligomer constructs, designed the coverslip adapter, developed staining protocols for antibody and *in situ* hybridization staining, directed the work, and wrote the paper. I.V. developed staining protocols, carried out the experimental work, and contributed to writing the paper. S.S.P. was engaged in discussion of tyramide-oligomer constructs, developing the staining protocols, and writing the paper.

