## Supplemental Information for "Indexable signal amplification for multiparametric imaging"

### Supplementary Methods

#### Institutional oversight

All protocols were carried out in accordance with relevant guidelines, regulations, and approval by the institutional review board at Weill Cornell Medicine (IRB #0107004999).

#### Validation of custom coverslip holders

Custom coverslip holders (see Fig. S1) were developed using FreeCAD software and were custom manufactured using 6061-T651 aluminum (ProtoLabs, Maple Plain, MN).

Staining of tonsil tissue on microscope slides and coverslips was performed for comparison using standard immunohistochemistry for CD163 as follows. Coverslips were charged by coating with poly-L-lysine per instructions for CODEX imaging (Akoya Biosciences CODEX Manual, revision C). 4-micron-thick sections of tonsil were then cut and placed on the charged coverslips or on charged microscope slides. The tissue sections were stained using immunohistochemistry with CD163 (clone 10D6, cat. no. PA0090) antibody in a Leica BondRX tissue staining instrument using standard immunohistochemical staining protocols outlined as follows: 1. Tissue was deparaffinized and rehydrated using ethanol washes. 2. Antigen retrieval was performed using an EDTA solution (Leica ER2 solution) at pH 9.0 at 95 degrees Celsius for 20 minutes. 3. Peroxidase activity was blocked with 3-4% hydrogen peroxide solution (Leica) for 5 minutes. 4. Tissue was stained with CD3e antibody (1:100 dilution, SP7 clone, Sigma, cat. no. SAB5500058) for 30 minutes at room temperature. 5. Secondary antibody staining using goat anti-rabbit antibodies conjugated to peroxidase (Leica) was performed (8 minutes, room temperature). 6. Incubation with DAB was performed to develop color. 7. Counterstaining was performed using hematoxylin. The resulting immunohistochemical staining highlighted cells consistent with histiocytes, as expected for CD163 staining. No qualitative difference in images obtained from tissue on coverslips or microscope slides was observed (see Fig. S2).

We next compared tonsil tissue stained on coverslips with tonsil tissue stained on microscope slides using tyramide-barcode staining and fluorescent complementary DNA oligomers (for protocol, see “Tyramide-barcode staining with a single antibody” below). We used CD3e primary antibody (clone SP7, 1:100 dilution, Sigma, cat. no. SAB5500058) for comparison between the two. Again, we saw no qualitative difference in staining quality for sections of tonsil placed on coverslips (using the custom coverslip holder for staining in a Leica Bond RX stainer) and microscope slides (see Fig. S3).

Of note, since the currently available commercial CODEX instrument does not allow the use of microscope slides, direct comparison between stained coverslips and stained microscope slides by CODEX imaging to evaluate the impact of the custom coverslip holder is not currently possible since, in the currently available instrument, microscope slides cannot be used.

### Tyramide-barcode staining with a single antibody

De-identified tonsil tissue was cut into 4-micron-thick sections and placed on charged coverslips or glass slides. The basic staining protocol for a single antibody using a Leica Bond RX autostainer is as follows, with Leica Bond Wash solution (Leica AR9590) used for washing where not otherwise stated: deparaffinization (Leica deparaffinization solution, cat. no. AR9222), washing (100% ethanol x3, Bond Wash solution x3), antigen retrieval (Leica ER2 solution for 20 minutes or ER1 solution for 30 minutes at 95 degrees Celsius), washing 5x, peroxide block (Leica peroxide block solution consisting of 3-4% hydrogen peroxide, 5 minutes), washing 3x, staining with primary antibody (30 minutes), washing 3x, staining with species-specific secondary antibody conjugated to peroxidase (8 minutes), washing 3x, incubating with 400 nM double-stranded tyramide-barcode conjugate in TE buffer (30 minutes), DAPI solution (Akoya Nuclear Stain, cat. no. FP1490, diluted 3 drops into 1 ml of 1x TBS) incubation for 5 minutes, washing 3x, and subsequent imaging on a CODEX instrument with appropriate single-stranded fluorescent complementary barcodes used for staining after tissue clearing (see Figs. 3-5). If cyclical imaging on a CODEX instrument was not to be used, the peroxidase was incubated with double-stranded barcode with attached fluorophore rather than non-fluorescent double-stranded tyramide-barcode conjugate (e.g., Fig. 2).

### Proof of principle using PAX-5 and CD19 co-staining in tonsil

De-identified tonsil tissue was cut into 4-micron-thick sections and placed on charged slides. Using a Leica BondRX automated stainer and our custom coverslip holder, the tissue was deparaffinized, followed by rehydration with ethanol washes and antigen retrieval (Leica ER2 solution, 20 minutes, 95˚C). The protocol then proceeded as follows with Bond Wash Solution (Leica) used for washing: Leica 3-4% hydrogen peroxide peroxidase blocking solution (5 minutes), wash 3x, mouse PAX-5 antibody (clone 24/PAX-5, BD Transduction Laboratories, cat. no. 610863) diluted 1:50 in Abcam antibody diluent (Abcam, cat. no. ab64211) for 30 minutes, wash 3x for 2 minutes each, Leica rabbit anti-mouse antibody (Leica) for 8 minutes, wash 3x (2 minutes each), Leica anti-rabbit antibody conjugated with HRP (8 minutes), wash 3x for 2 minutes each, 500 nM double-stranded tyramide-barcode (designated #79) with Cy3 on the reporter strand in TE buffer (30 minutes), wash (5 minutes), Leica 3-4% hydrogen peroxide peroxidase blocking solution (5 minutes), wash 3x, CD19 (BT51E, Leica cat.no. PA0843) ready-to-use antibody solution (30 minutes), wash 3x for 2 minutes each, Leica rabbit anti-mouse antibody (8 minutes), wash 3x for 2 minutes each, Leica anti-rabbit antibody conjugated with HRP (8 minutes), wash 3x for 2 minutes each, 500 nM double-stranded tyramide-barcode (designated #77; for corresponding DNA sequence, see Schurch et al.[1]) with Cy5 on the reporter strand in TE buffer (30 minutes), and de-ionized water 2x for 10 minutes each. Images were captured with a Keyence BZ-X800 microscope with 20x Nikon PlanApo 0.75 NA objective and "high resolution" camera settings with ATTO550/Cy3 and Cy5 filter sets.

Results demonstrated that both PAX-5 and CD19 antibodies appropriately stained cells consistent with B cells, with PAX-5 antibody staining B cell nuclei and CD19 antibody staining B cell membranes (see Fig. S4).

### Comparison of tyramide-barcodes and tyramide-fluorophores via PAX-5 and CD19 co-staining in tonsil

For comparison to verify consistent fluorophore localization and to rule out unforeseeable effects due to the DNA barcodes, side-by-side staining with tyramide-barcodes with attached fluorescent complementary reporter strands versus tyramide conjugated directly to fluorophores, a well established technique, was performed.

Staining was performed as follows in a Leica BondRX autostainer: Deparaffinize tonsil tissue placed on charged slides, wash (100% ethanol x3, Bond Wash solution x3), antigen retrieval using Leica ER2 solution for 20 minutes at 95˚C, Leica 3-4% hydrogen peroxide peroxidase blocking solution (5 minutes), wash 3x with Leica Bond Wash solution (0 minutes per wash), mouse PAX-5 antibody (clone 24/PAX-5, BD Transduction Laboratories, cat. no. 610863) diluted 1:50

in Abcam antibody diluent (Abcam, cat. no. ab64211) for 30 minutes, wash 3x with Leica Bond Wash solution (2 minutes per wash), Leica rabbit anti-mouse secondary antibody ready-to-use solution (Leica, 8 minutes), wash 3x with Leica Bond Wash solution (2 minutes per wash), Leica anti-rabbit tertiary antibody HRP conjugate ready-to-use solution (8 minutes), wash 3x with Leica Bond Wash solution (2 minutes per wash), 500 nM double-stranded tyramide-barcode79 with Cy3 on the reporter strand in TE buffer (30 minutes) *or* 500 nM tyramide-Cy3 (R&D Systems, cat. no. 6457/1, diluted to 10 mM stock solution in DMSO followed by final dilution to 500 nM in TE buffer) for 30 minutes, wash with Leica Bond Wash solution (5 minutes), incubate with Leica 3-4% hydrogen peroxide peroxidase blocking solution (5 minutes), wash 3x with Leica Bond Wash solution (2 minutes per wash), incubate with CD19 (clone BT51E, Leica cat. no. PA0843) ready-to-use solution for 30 minutes, wash 3x with Leica Bond Wash solution (2 minutes per wash), incubate with Leica rabbit anti-mouse secondary antibody ready-to-use solution (8 minutes), wash 3x with Leica Bond Wash solution (2 minutes per wash), incubate with Leica anti-rabbit tertiary antibody HRP conjugate solution (8 minutes), wash 3x with Leica Bond Wash solution (2 minutes per wash), incubate with 500 nM double-stranded tyramide-barcode77 with Cy5 on the reporter strand in TE buffer *or* 500 nM tyramide-Cy5 (R&D Systems, cat. no. 6458/1, diluted to 10 mM stock solution in DMSO followed by final dilution to 500 nM in TE buffer) for 30 minutes, wash 3x with Leica Bond Wash solution, DAPI solution (Akoya Nuclear Stain, 3 drops per mL) incubation for 5 minutes, wash 3x with Leica Bond Wash solution.

Images were captured with a Keyence BZ-X800 microscope with 20x Nikon PlanApo 0.75 NA objective and "high resolution" camera settings with DAPI, ATTO550/Cy3, and Cy5 filter sets. No significant difference in distribution of staining was seen when compared using PAX-5 and CD19 primary antibodies, which stain B cells (see Fig. S5).

### Comparison of staining intensity by barcode-conjugated primary antibodies and tyramide-barcodes

In brief, two tonsil tissue sections were stained by barcode-conjugated antibodies. One was further stained with tyramide barcodes. Both tissue sections were imaged using CODEX imaging, using Cy5-conjugated complementary oligomers appropriate to each.

In more detail, the CODEX staining using standard barcode-conjugated primary antibodies was performed as follows: In a Leica BondRX autostainer, deparaffinize tonsil tissue that is placed on a charged coverslip, wash (100% ethanol x3, Leica Bond Wash solution x3), antigen retrieval (Leica ER2 solution, EDTA pH 9.0) for 20 minutes at 95˚C, wash (Leica Bond Wash solution), Akoya hydration buffer (2 min. x2), Akoya Staining Buffer (20 minutes), and staining with CD3e (1:200 dilution, Akoya, cat. no. 4450030) primary antibody diluted in Abcam antibody dilution solution (Abcam, cat. no. ab64211) for 90 minutes x2. Remove sample from autostainer. Incubate the coverslip in Akoya Staining Buffer (in a well of a tissue culture plate) for 2 minutes, incubate the coverslip in Akoya Staining Buffer (a second well of a tissue culture plate) for 2 minutes, incubate the coverslip in the post-staining fixation solution (0.5 ml 16% paraformaldehyde [PFA, Alfa Aesar, cat. No 43368] in 4.5 ml Akoya Staining Buffer) for 10 minutes, wash 3x with PBS, incubate the coverslip in ice cold methanol for 5 minutes, and wash 3x with PBS. Image using an Akoya CODEX instrument paired with a Keyence BZ-X800 fluorescence microscope with 20x Nikon PlanApo lambda objective per Akoya CODEX Manual (revision C), using Cy5-conjugated complementary barcode strands and 500 ms acquisition time.

Staining using tyramide-barcodes was performed as follows: In a Leica BondRX autostainer, deparaffinize tonsil tissue that is placed on a charged coverslip, wash (100% ethanol x3, Leica Bond Wash solution x3), antigen retrieval (Leica ER2 solution, EDTA pH 9.0) for 20 minutes at 95˚C, wash (Leica Bond Wash solution), Akoya hydration buffer (2 min. x2), Akoya Staining Buffer (20 minutes), staining with CD3e (1:200 dilution, Akoya, cat. no. 4450030) primary antibody diluted in Abcam antibody dilution solution (Abcam, cat. no. ab64211) for 90 minutes x2, Leica 3-4% hydrogen peroxide peroxidase blocking solution (5 minutes), wash 3x with Leica Bond Wash solution (2 minutes per wash), goat anti-rabbit secondary antibody HRP conjugate ready-to-use solution (Leica) (8 minutes), wash 3x with Leica Bond Wash solution (2 minutes per wash), and incubation with 500 nM tyramide-barcode (designated #77) and non-fluorescent complementary strand in TE buffer pH 8.0 (15 minutes). Remove sample from autostainer. Incubate the coverslip in

Akoya Staining Buffer (in a well of a tissue culture plate) for 2 minutes, incubate the coverslip in Akoya Staining Buffer (a second well of a tissue culture plate) for 2 minutes, incubate the coverslip in the post-staining fixation solution (0.5 ml 16% PFA [Alfa Aesar, cat. No 43368] in 4.5 ml Akoya Staining Buffer) for 10 minutes, wash 3x with PBS, incubate the coverslip in ice cold methanol for 5 minutes, and wash 3x with PBS. Image using an Akoya CODEX instrument paired with a Keyence BZ-X800 fluorescence microscope with 20x Nikon PlanApo lambda 0.75 NA objective per Akoya CODEX Manual (revision C), using 400 nM Cy5-conjugated complementary barcode strands in reporter buffer (see Akoya CODEX Manual, revision C) and 500 ms acquisition time.

Comparison of the staining intensities was performed using acquired raw TIFF files (not modified by post-imaging processing) using ImageJ. Histograms of the pixel intensity values were calculated for similar appearing tonsil tissue. Approximate modal values and half-modal values for non-background pixels (background pixels form a sharp peak in both histograms near zero) were calculated for both experiments, with the tyramide-barcodes experiment having approximately 4.7x greater modal and half-modal intensity values (see Figure S6).

### Comparison of staining using single-stranded versus double-stranded tyramide barcodes

Early failed experiments suggested that staining with single-stranded tyramide-barcodes was inferior to staining using double-stranded tyramide-barcodes. The reason is unclear as we were later unable to reproduce the failed experiment, but considerations include human error and possible contamination of solutions with DNAse. However, later experiments using single-stranded tyramide barcodes demonstrated consistent staining similar to that of double-stranded tyramide-barcode staining (see Figs. S7, S8, and S9). The protocols used for comparing single-stranded tyramide-barcodes and double-stranded tyramide-barcodes are as follows:

*Single-stranded tyramide-barcodes*. In a Leica BondRX autostainer, deparaffinize tonsil tissue that is placed on a charged coverslip, wash (100% ethanol x3, Leica Bond Wash solution x3), perform antigen retrieval (Leica ER2 solution, EDTA pH 9.0) for 20 minutes at 95˚C, wash 5x with Leica Bond Wash solution, incubate with Leica 3-4% hydrogen peroxide peroxidase blocking solution (5 minutes), wash 3x with Leica Bond Wash solution (0 minutes per wash), stain with CD3e (1;200 dilution, clone , Abcam, cat. no. ab52959) primary antibody diluted in Abcam antibody diluent (Abcam, cat. no. ab64211) for 30 minutes, wash 3x with Leica Bond Wash solution (2 minutes per wash), incubate with goat anti-rabbit secondary antibody HRP conjugate ready-to-use solution (Leica, cat. no. PV6119) (8 minutes), wash 3x with Leica Bond Wash solution (2 minutes per wash), incubate with 500 nM tyramide-barcode in TE buffer pH 8.0 (15 minutes), wash 3x with Leica Bond Wash solution (2 minutes per wash), incubate with 500 nM complementary barcode conjugated to Cy5 in TE buffer pH 8.0 (15 minutes), wash 3x with Leica Bond Wash solution (2 minutes per wash), incubate in DAPI solution (Akoya, cat. no. FP1490) diluted 3 drops per 1 mL of TBS pH 8.0 for 5 minutes, wash 1x with Leica Bond Wash solution (2 minutes per wash), collect images.

*Double stranded tyramide-barcodes*. In a Leica BondRX autostainer, deparaffinize tonsil tissue that is placed on a charged coverslip, wash (100% ethanol x3, Leica Bond Wash solution x3), antigen retrieval (Leica ER2 solution, EDTA pH 9.0) for 20 minutes at 95˚C, wash 5x with Leica Bond Wash solution, Leica 3-4% hydrogen peroxide peroxidase blocking solution (5 minutes), wash 3x with Leica Bond Wash solution (0 minutes per wash), staining with CD3e (1;200 dilution, clone , Abcam, cat. no. ab52959) primary antibody diluted in Abcam antibody diluent (Abcam, cat. no. ab64211) for 30 minutes, wash 3x with Leica Bond Wash solution (2 minutes per wash), incubate with goat anti-rabbit secondary antibody HRP conjugate ready-to-use solution (Leica, cat. no. PV6119) (8 minutes), wash 3x with Leica Bond Wash solution (2 minutes per wash), incubation with 500 nM tyramide-barcode and 500 nM tyramide-barcode in TE buffer pH 8.0 (15 minutes), incubation in DAPI solution (Akoya, cat. no. FP1490) diluted 3 drops per 1 mL of TBS pH 8.0 for 5 minutes, wash 1x with Leica Bond Wash solution (2 minutes per wash), collect images.

*Imaging*. Image on Keyence BZ-X800 fluorescence microscope ("high-resolution" mode) with 20x Nikon PlanApo lambda objective and Cy5 filter set, 0.5 s acquisition times.

### EBER in situ hybridization staining using tyramide-barcodes

4-micron-thick sections of EBV+ classic Hodgkin lymphoma tissue are placed on charged coverslips or glass slides. The staining protocol is as follows: deparaffinization (30 minutes at 72°), 100% ethanol wash x3, proteinase K enzyme digestion (13 minutes at 37°C), washing 4x, incubate with FITC-conjugated EBER probe for 60 minutes, wash 15x, incubate with Leica ready-to-use mouse anti-FITC antibody (15 minutes), wash 3x, incubate with Leica ready-to-use rabbit anti-mouse antibody (10 minutes), wash 3x (2 minutes per wash), incubate with Leica ready-to-use peroxidase-conjugated anti-rabbit antibody (15 minutes), wash x9, incubate with ~0.1 mg/mL double-stranded tyramide-barcode conjugate in TE buffer (20 minutes), wash, incubate in DAPI solution (Akoya, cat. no. FP1490) diluted 3 drops per 1 mL of TBS, wash x3.

### Multiplexed tyramide-barcode staining with cyclical imaging

Tissue sections were placed on charged coverslips and stained using the custom coverslip holder described above. After deparaffinization, antigen retrieval (within the Leica Bond RX autostainer), and washing, as described for protocols above, an antibody staining cycle was performed for each one of the antibodies used.

For mouse primary antibodies, the staining cycle protocol consisted of washing (with Leica Bond Wash solution unless stated otherwise), peroxidase block in Leica 3-4% hydrogen peroxide solution (5 minutes), washing 3x, incubation with primary antibody (30 minutes), washing 2x (2 minutes each), incubation with Leica rabbit anti-mouse antibody ready-to-use solution (8 minutes, Leica), washing 3x (2 minutes each), incubation with Leica ready-to-use peroxidase-conjugated anti-rabbit antibody (8 minutes, Leica), washing 3x (2 minutes each), incubation with 1 mg/15 ml double-stranded tyramide barcode in TE solution (30 minutes), washing at 95˚C for 5 minutes, washing 2x, incubation with Leica Bond Wash at 95˚C for 20 minutes, and washing.

For rabbit primary antibodies, the staining cycle protocol consisted of washing, peroxidase block in Leica 3-4% hydrogen peroxide solution (5 minutes), washing 3x, incubation with primary antibody (30 minutes), washing 3x (2 minutes each), incubation with Leica anti-rabbit antibody ready-to-use solution (8 minutes, Leica), washing 3x (2 minutes each), 1 mg/15 ml double-stranded tyramide barcode (30 minutes), washing at 95˚C for 5 minutes, washing 2x, washing at 95˚C for 20 minutes, and washing.

After all antibody staining cycles were complete, the ending protocol consisted of washing at 95˚C for 5 minutes, incubation in DAPI solution (Akoya, cat. no. FP1490) diluted 3 drops per 1 mL of TBS, and washing again at 95˚C for 5 minutes.

### Supplementary Figures

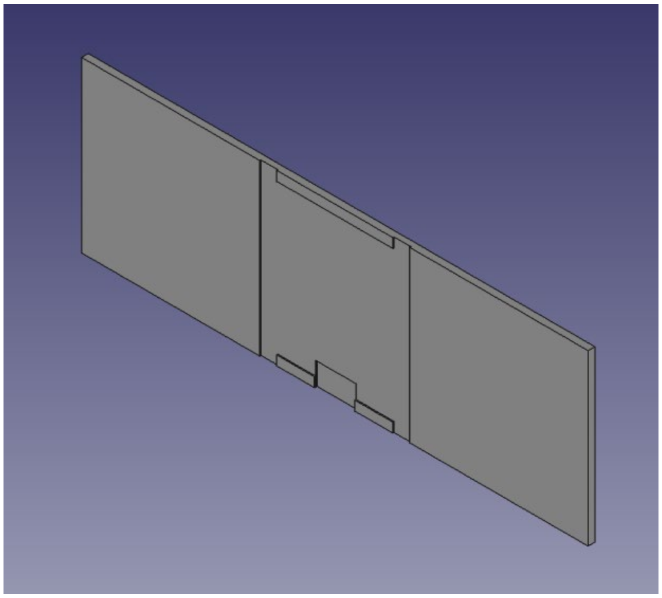

Figure S1. Coverslip holder design for staining coverslips on a Leica Bond RX tissue autostainer. The slots at the four corners of the coverslip placement site are in place to facilitate manufacturing and ensure a good fit for the coverslip. The pocket on the side (bottom edge) facilitates lifting the coverslip out of the holder.

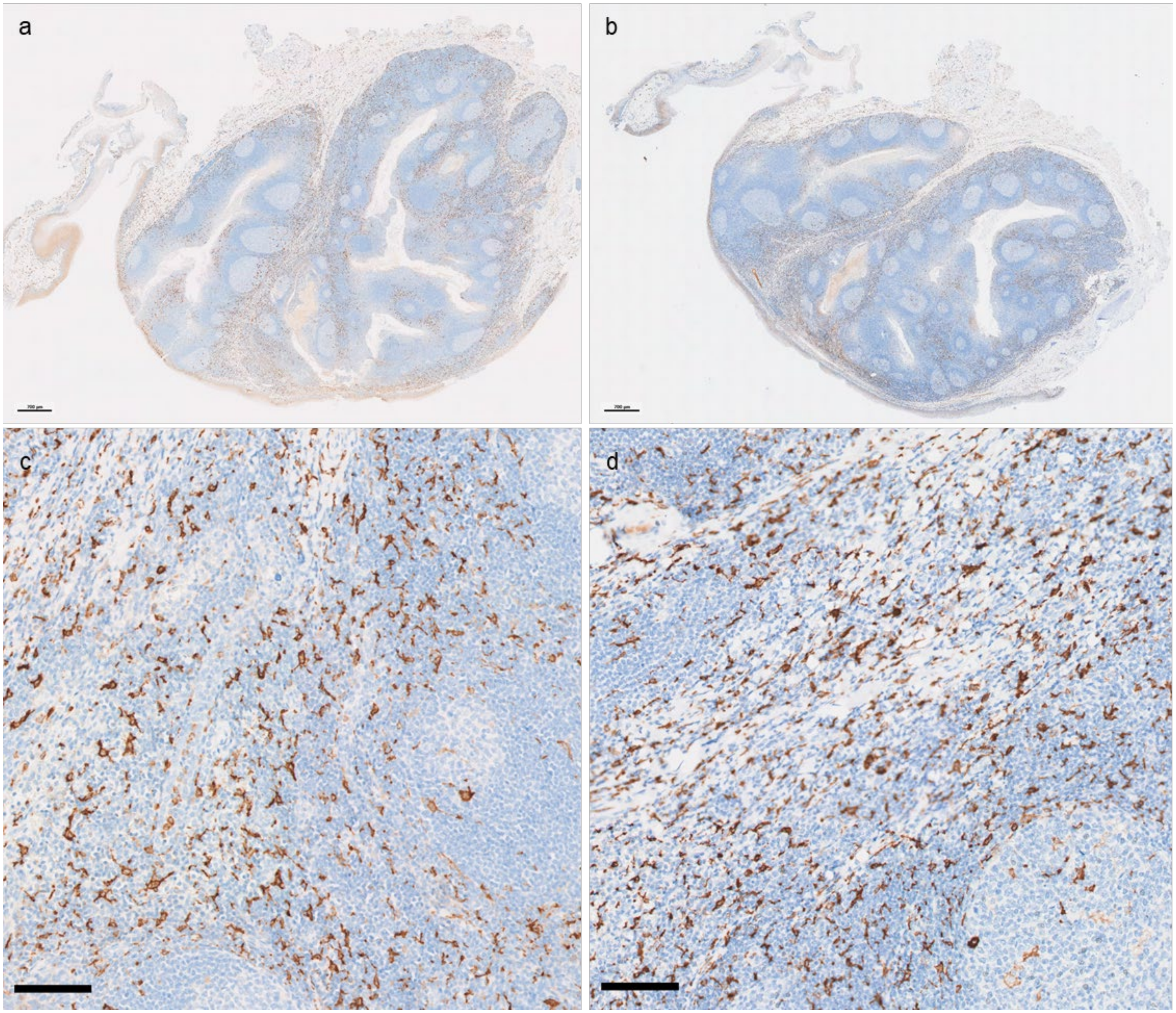


Figure S2. Comparison of CD163 IHC staining obtained using a glass slide (**a**, **c**) and a coverslip stained using the custom coverslip holder (**b**, **d**) and otherwise identical staining protocols. No significant qualitative difference in staining quality was identified. Slides were scanned using a Perkin-Elmer Vectra Polaris instrument at 20x magnification. Scale bars in **a** and **b** are 700 microns. Scale bars in **c** and **d** are 100 microns.

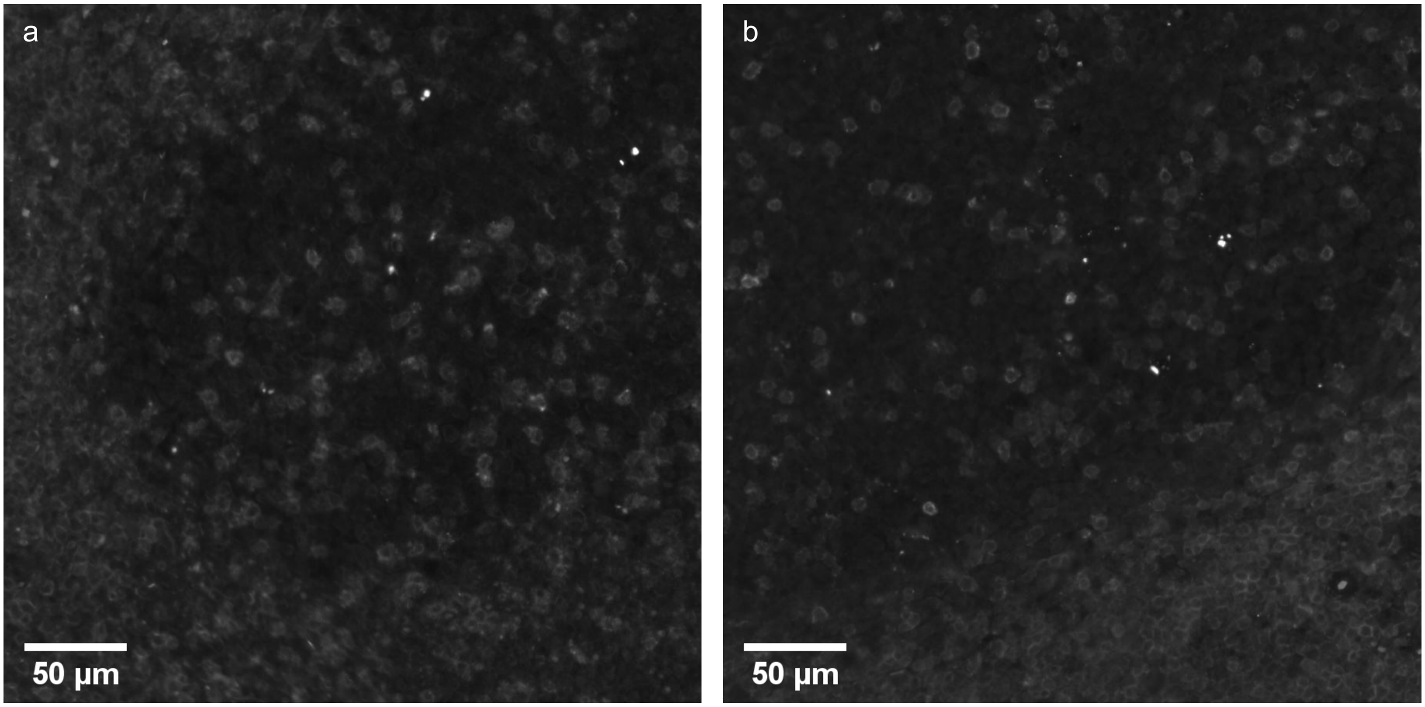


Figure S3. Comparison of tonsil tissue on a microscope slide versus a coverslip stained with CD3e primary antibody (clone SP7, Sigma, cat. no. SAB5500058-100UL) and tyramide-barcode staining. Tonsil tissue was placed on a charged microscope slide (**a**) and a charged coverslip (**b**), which then were deparaffinized, washed (100% ethanol x3, Bond Wash solution x3), underwent antigen retrieval (Leica ER2 solution, EDTA pH 9.0) for 20 minutes at 95˚, washed 5x with Leica Bond Wash solution, incubated with Leica 3-4% hydrogen peroxide peroxidase blocking solution (5 minutes), washed 3x with Leica Bond Wash solution (0 minutes per wash), stained with CD3e (1:100 dilution, clone SP7, Sigma, cat. no. SAB5500058-100UL) primary antibody diluted in Abcam antibody diluent (Abcam, cat. no. ab64211) for 30 minutes, washed 3x with Leica Bond Wash solution (2 minutes per wash), incubated with goat anti-rabbit secondary antibody and peroxidase conjugate ready-to-use solution (Leica, 8 minutes), washed 3x with Leica Bond Wash solution (2 minutes per wash), incubated with 500 nM tyramide-barcode77 with Cy5-conjugated reporter strand in TE buffer (15 minutes), incubated 5 minutes in DAPI solution (Akoya Nuclear Stain, diluted 3 drops per 1 mL of TBS), washed 1x with Leica Bond Wash solution (2 minutes), and imaged on a Keyence BZ-X800 fluorescence microscope with 20x Nikon PlanApo lambda objective and DAPI and Cy5 appropriate filter sets. The images show no significant difference in intensity or distribution of staining when comparing the microscope slide (**a**) and coverslip (**b**).

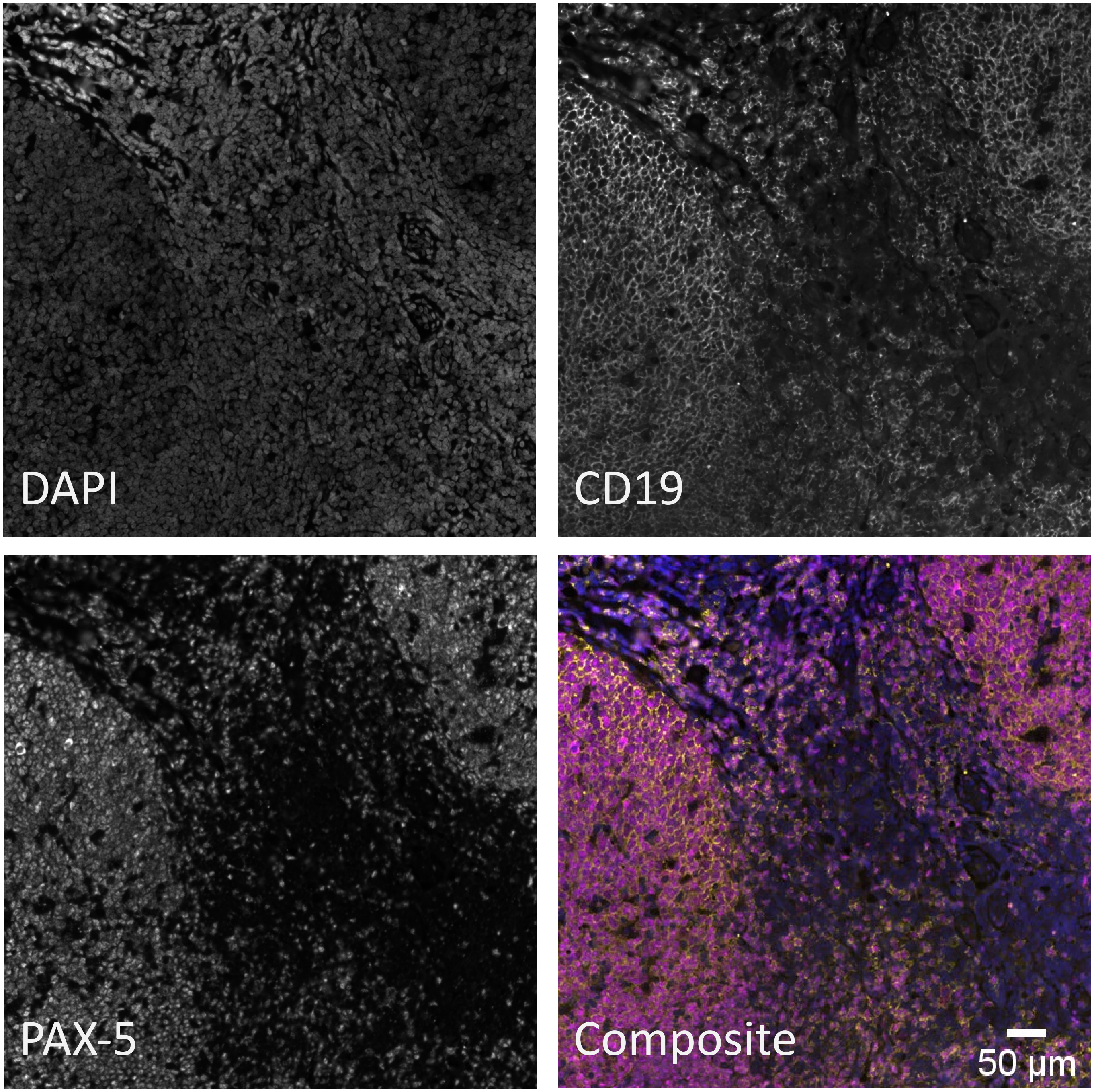


Figure S4. Multiplexed tyramide-barcode staining of tonsil. Tonsil was stained on a Leica Bond RX autostainer with DAPI nuclear stain, mouse anti-CD19 antibody (ready-to-use solution, clone BT51E, Leica cat. no. PA0843), and mouse anti-PAX-5 antibody (1:50 dilution, clone 24/PAX-5, BD Transduction Laboratories, cat. no. 610863) primary antibodies (in different staining steps). The primary antibodies were followed by Leica rabbit anti-mouse antibodies (ready-to-use solution) and peroxidase conjugated tertiary antibodies (ready-to-use solution, Leica), which were subsequently allowed to react with tyramide-barcode-Cy3 (for CD19) and tyramide-barcode-Cy5 (for PAX-5). The composite image shows DAPI-stained nuclei in blue, CD19-stained cell membranes in yellow, and PAX-5 stained nuclei in magenta and demonstrates the appropriate co-localization of PAX-5 and CD19 in B cells. Images were captured with a Keyence BZ-X800 microscope with 20x Nikon PlanApo 0.75 NA objective and "high resolution" camera settings. 50 µm scale bar applies to all 4 panels.

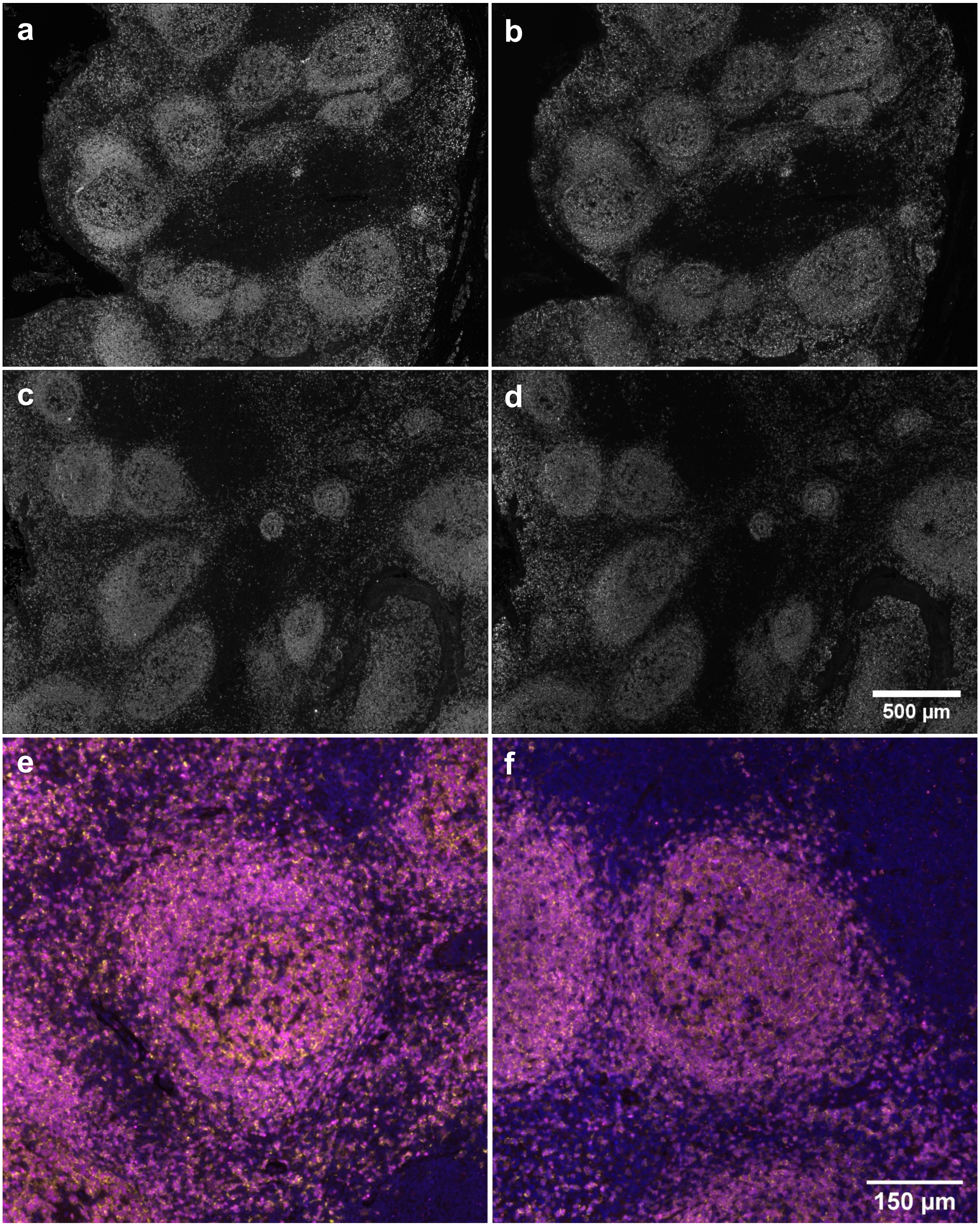


Figure S5. Comparison of staining with tyramide-barcodes with fluorescent complementary oligomers versus staining with tyramide conjugated directly to fluorophores. Parts **a** and **b** demonstrate tonsil stained with *tyramide-barcodes*, with PAX-5 highlighted with Cy3-conjugated complementary oligomer in **a**, and CD19 highlighted with Cy5-conjugated complementary oligomer in **b**. Parts **c** and **d** demonstrate tonsil stained with *tyramide conjugated directly to fluorophores*, with PAX-5 highlighted with Cy3 in **a**, and CD19 highlighted with Cy5 in **d**. In all four parts (**a**-**d**), staining is consistent with staining of B cells, found primarily in germinal centers and associated mantle and marginal zones. Part **e** demonstrates a composite image of tyramide barcode staining (parts **a** and **b**) plus DAPI staining. Part **f** demonstrates a composite image of tyramide-fluorophore staining (parts **c** and **d**), plus DAPI. The composite images show DAPI-stained nuclei in blue, CD19-stained cell membranes in yellow, and PAX-5 stained nuclei in magenta and demonstrate the appropriate co-localization of PAX-5 and CD19 in B cells. Images were captured with a Keyence BZ-X800 microscope with 20x Nikon PlanApo 0.75 NA objective and "high resolution" camera settings and equivalent acquisition times for both tyramide-barcode and tyramide-fluorophore stains. Scale bar in **d** applies to **a-d**. Scale bar in **f** applies to **e** and **f**.

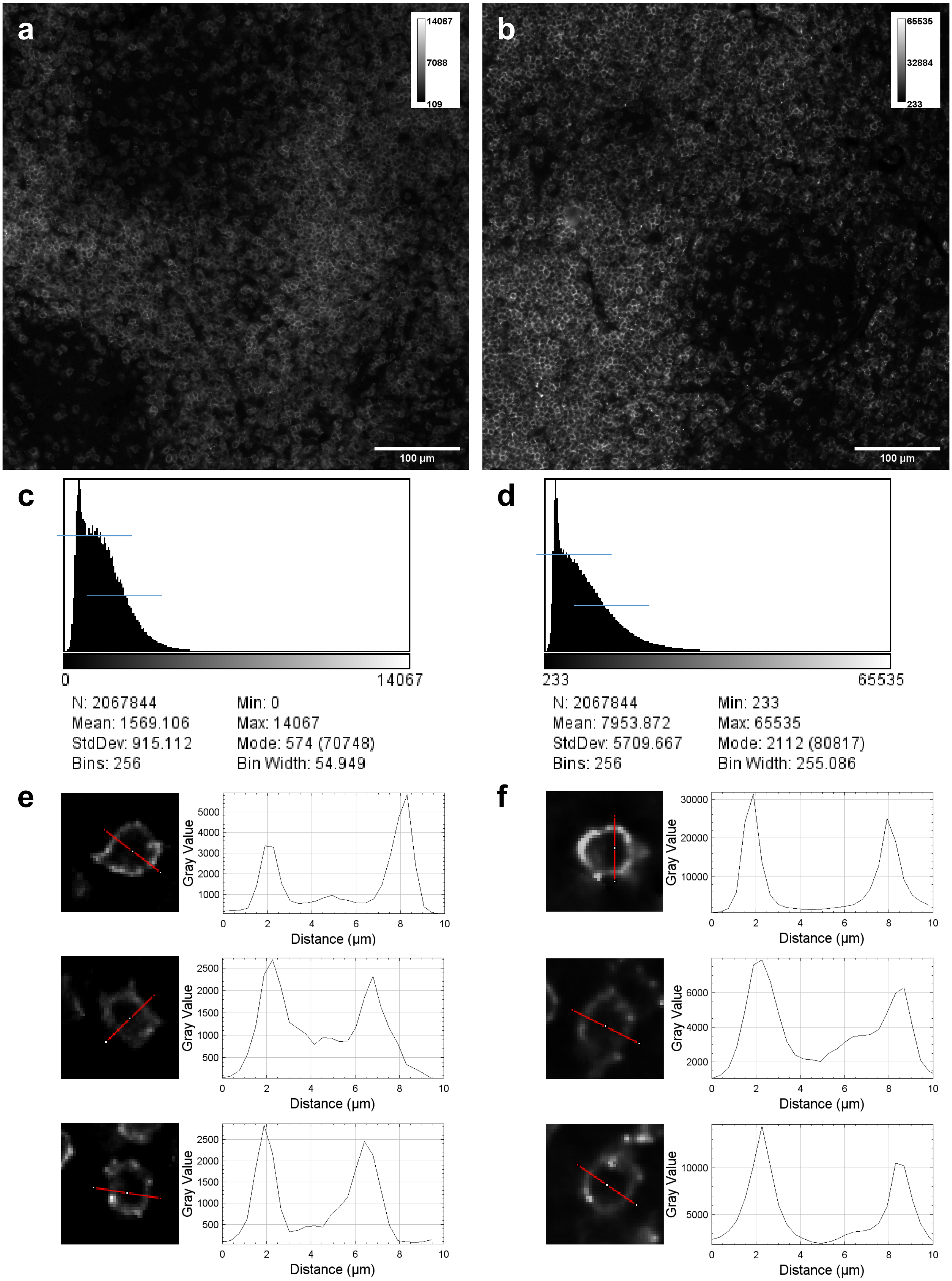

a
14067
7088
109
100 µm
b
65535
32884
233
100 µm
c
0
14067
N: 2067844
Mean: 1569.106
StdDev: 915.112
Bins: 256
Min: 0
Max: 14067
Mode: 574 (70748)
Bin Width: 54.949
d
233
65535
N: 2067844
Mean: 7953.872
StdDev: 5709.667
Bins: 256
Min: 233
Max: 65535
Mode: 2112 (80817)
Bin Width: 255.086
e
f
Gray Value
Distance (µm)

Figure S6. Comparison of tissue stained by conventional CODEX staining and tyramide-barcode CODEX staining. To verify that the method actually amplifies signal, tonsil tissue was stained with barcode-conjugated antibody for CD3e (1:200 dilution, Akoya Biosciences, cat. no. 4450030). This was followed by CODEX imaging using the corresponding Cy5-conjugated complementary DNA oligomer (panel **a**). For comparison, a section of the same tissue was stained with the same primary antibody, but then followed by incubation with secondary antibody with attached peroxidase, then incubation with tyramide-conjugated barcode, followed by CODEX imaging using Cy5-conjugated complementary oligomer specific for the tyramide-barcode (panel **b**). Panels **a** and **b** demonstrate appropriate localization of staining to T cells in the images, albeit with very different fluorescence intensity distributions, which intensity distributions are also shown in parts **c** and **d** (note the scale bars of the plots). Comparison of approximate modal values and half modal values (approximately indicated by blue lines shown in **c** and **d**), demonstrates approximately 4.7-fold increase in staining intensity for the tyramide-barcode staining over regular CODEX staining. Parts **e** and f demonstrate cross sectional image intensity profiles for a small sample of individual cells to get a sense of imaging resolution achieved using conventional CODEX imaging (**e**) and tyramide-barcodes imaging (**f**); the imaging resolutions appear similar for the given experiments. Images were captured using an Akoya CODEX instrument and equivalent camera settings for both experiments on a Keyence BZ-X800 microscope with 20x Nikon PlanApo 0.75 NA objective.

Barcode Single-stranded Double-stranded

72

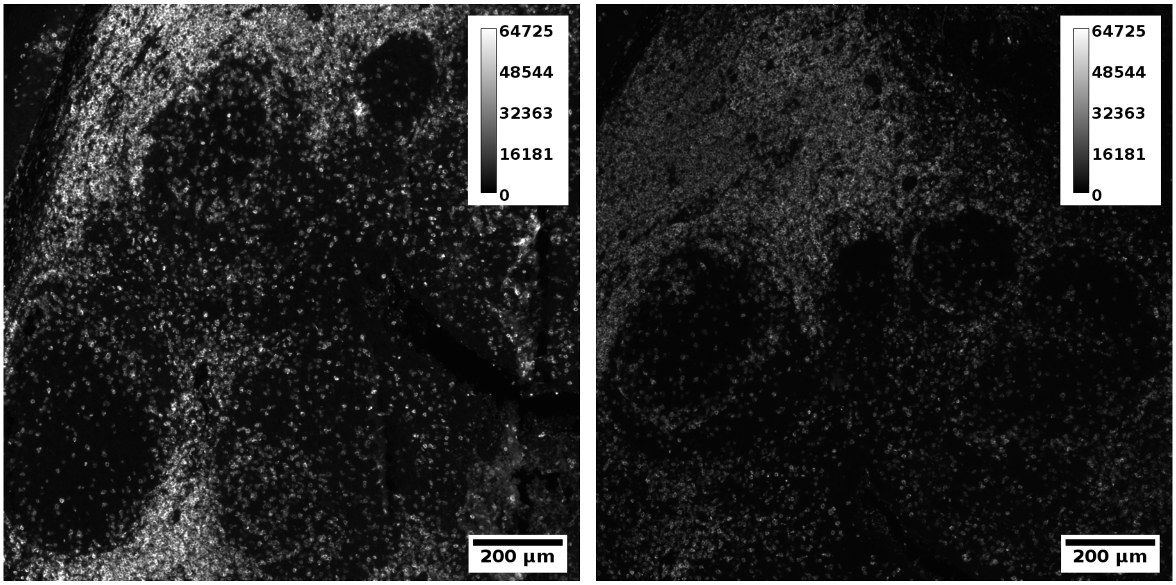


Figure S7. Comparison of CD3e staining using single-stranded versus double-stranded tyramide-barcodes in the step involving incubation with peroxidase demonstrates that single-stranded tyramide barcodes and double-stranded tyramide barcodes perform similarly well. In this figure, results are shown for barcode “72”. Comparisons for barcodes “74”, “75”, “76”, “77”, “79”, and “80” are shown in Figs. S8 and S9. All images were recorded using the same camera settings, including 0.5 s camera acquisition times. See also the experiment details in Supplementary Methods. Note that our early experiments failed to demonstrate staining with single-stranded tyramide barcodes for unknown reasons; however, the findings could not be replicated as later experiments all successfully demonstrated staining with single-stranded DNA barcodes.

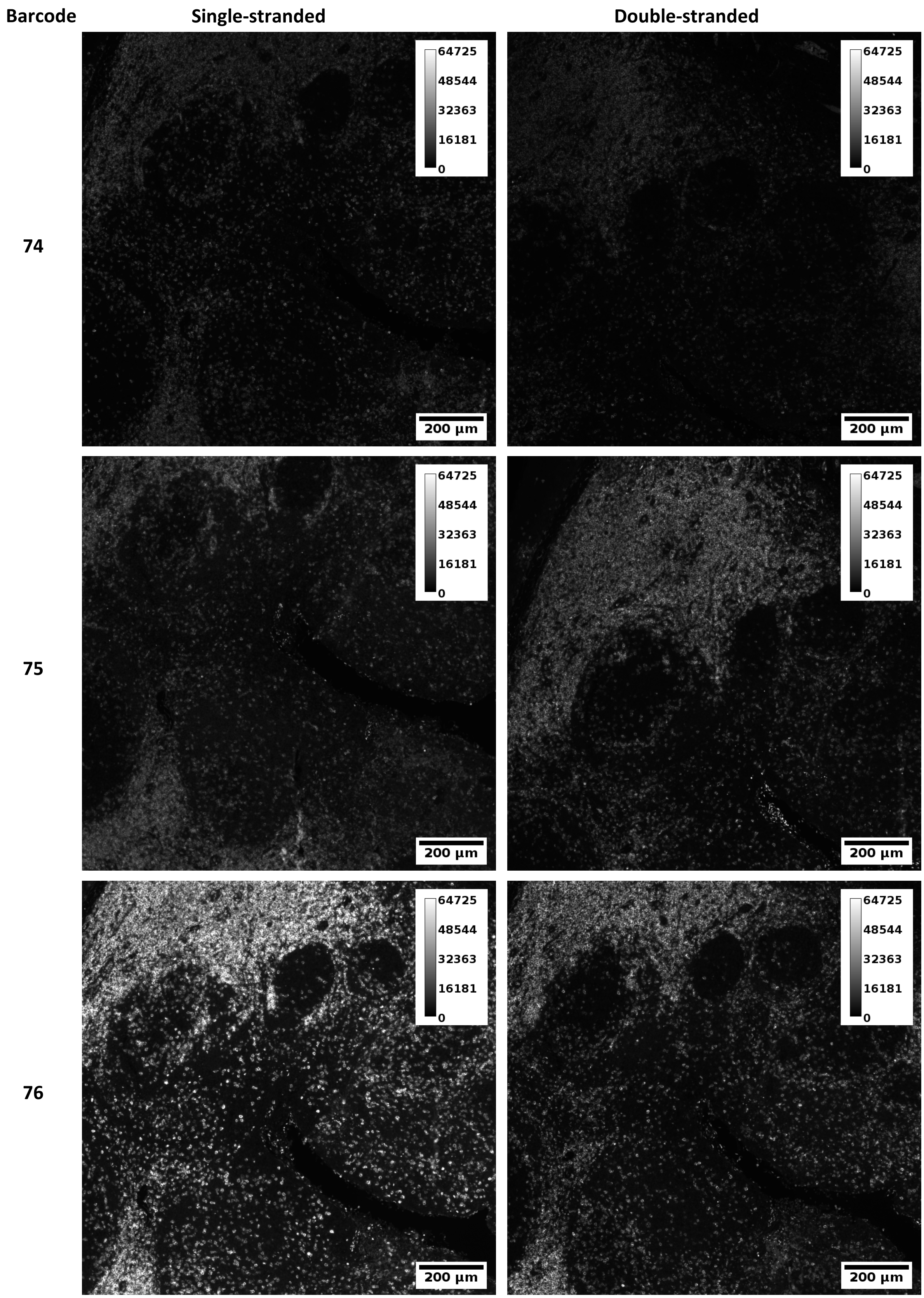


Figure S8. Comparison of single-stranded versus double-stranded tyramide-barcode staining for barcodes labeled “74”, “75”, and “76”. See also Figs. S7 and S9.

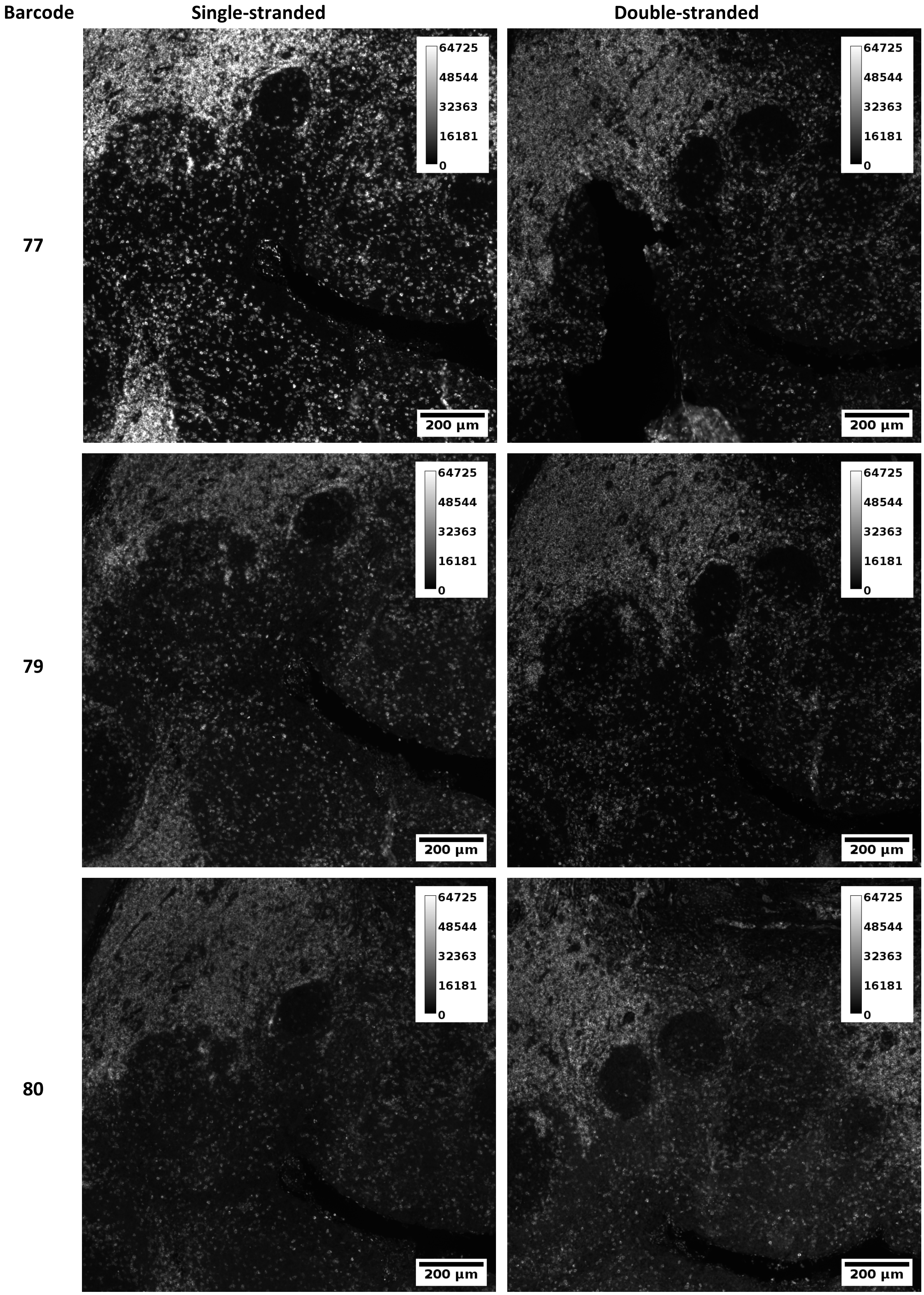


Figure S9. Comparison of single-stranded versus double-stranded tyramide-barcode staining for barcodes labeled “77”, “79”, and “80”. See also Figs. S7 and S9.

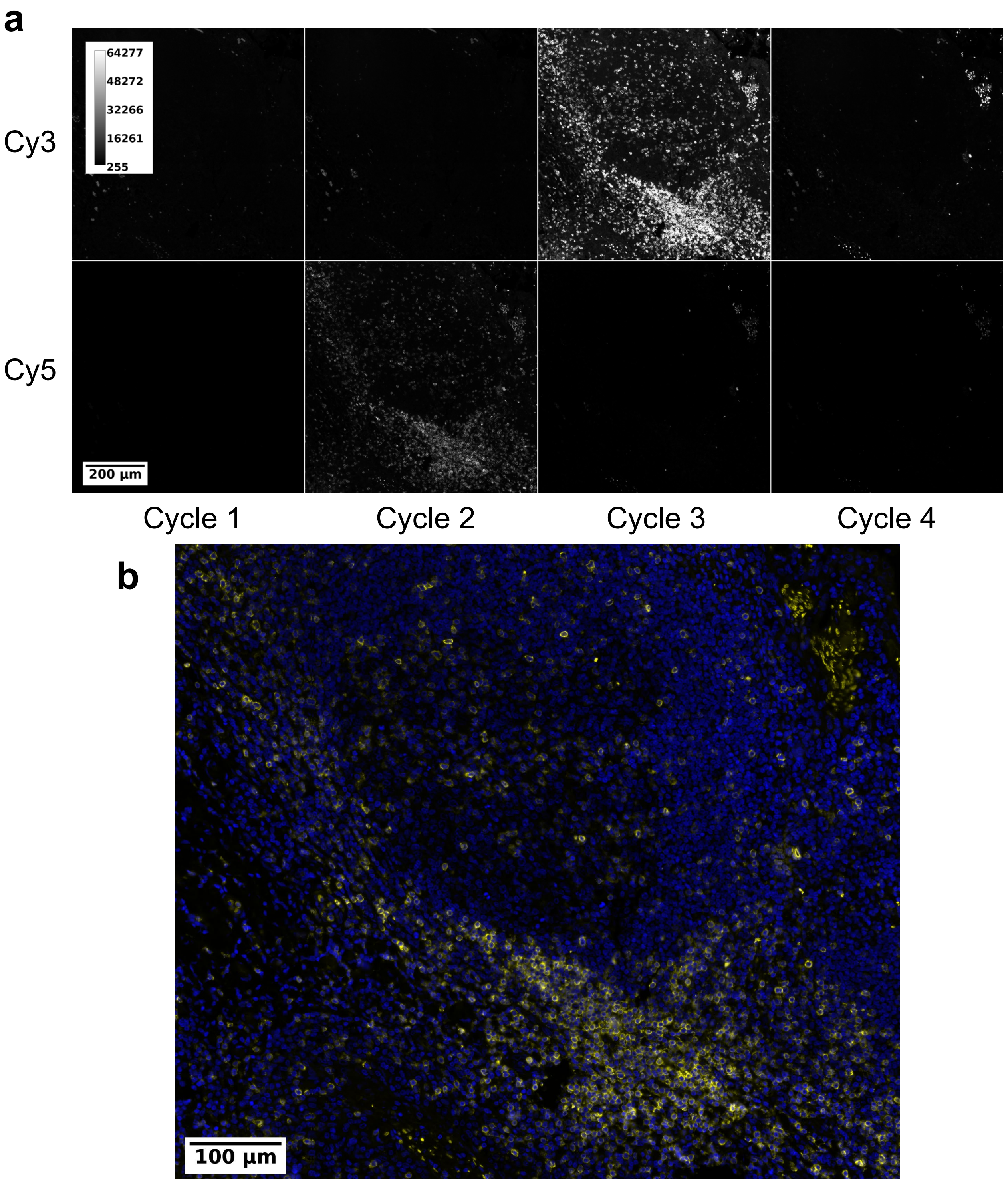


Figure S10. Demonstration of appropriate cycling with tyramide-barcode stained tonsil using an Akoya CODEX instrument and Keyence BZ-X800 microscope. Prior to imaging, tissue was stained using a Leica Bond RX autostainer with rabbit anti-CD3e primary antibody, peroxidase-conjugated secondary antibody, and tyramide-barcode. The tissue was then imaged on the CODEX/Keyence system using four imaging cycles; the tissue is cleared between each cycle. (**a**) In cycle 1, no complementary barcode was added; hence no staining was seen. In cycle 2, Cy5-conjugated reporter (complementary barcode) was added, with appropriate staining seen in the Cy5 imaging channel. In cycle 3, Cy3-

conjugated reporter was added, with appropriate imaging seen in the Cy3 channel. In cycle 4, no reporter was added, and appropriate clearing of reporters is seen. DAPI staining was performed but is not shown. Though appropriate clearing is seen, close inspection demonstrates very weak persistent signal in the Cy3 channel in Cycle 4 and in the Cy5 channel for Cycle 3, both demonstrating approximately 70-fold decrease from the cycles immediately prior. Cycle 4 of the Cy5 channel demonstrates a further 2-fold decrease in the residual signal, attributed to either photobleaching or further clearing of the reporter strands by the CODEX instrument. Intensity and size scales apply to all panes of part **a**. (**b**) False-colored image consists of DAPI (blue) and CD3e tyramide-barcode staining with Cy5-conjugated reporter (yellow). DAPI staining helps to demonstrate a germinal center while CD3e staining highlights T lymphocytes surrounding and scattered within the germinal center. Camera/microscope settings: 200 ms exposure time in Cy3 and Cy5 channels, 100% excitation intensity, "high resolution" imaging, Nikon 20x PlanApo lambda 0.75 NA objective. Camera pixels were saturated by the Cy3 intensity in cycle 3, demonstrating a need to further reduce the exposure time. Background subtraction and image stitching were performed using Akoya's CODEX Processor software.

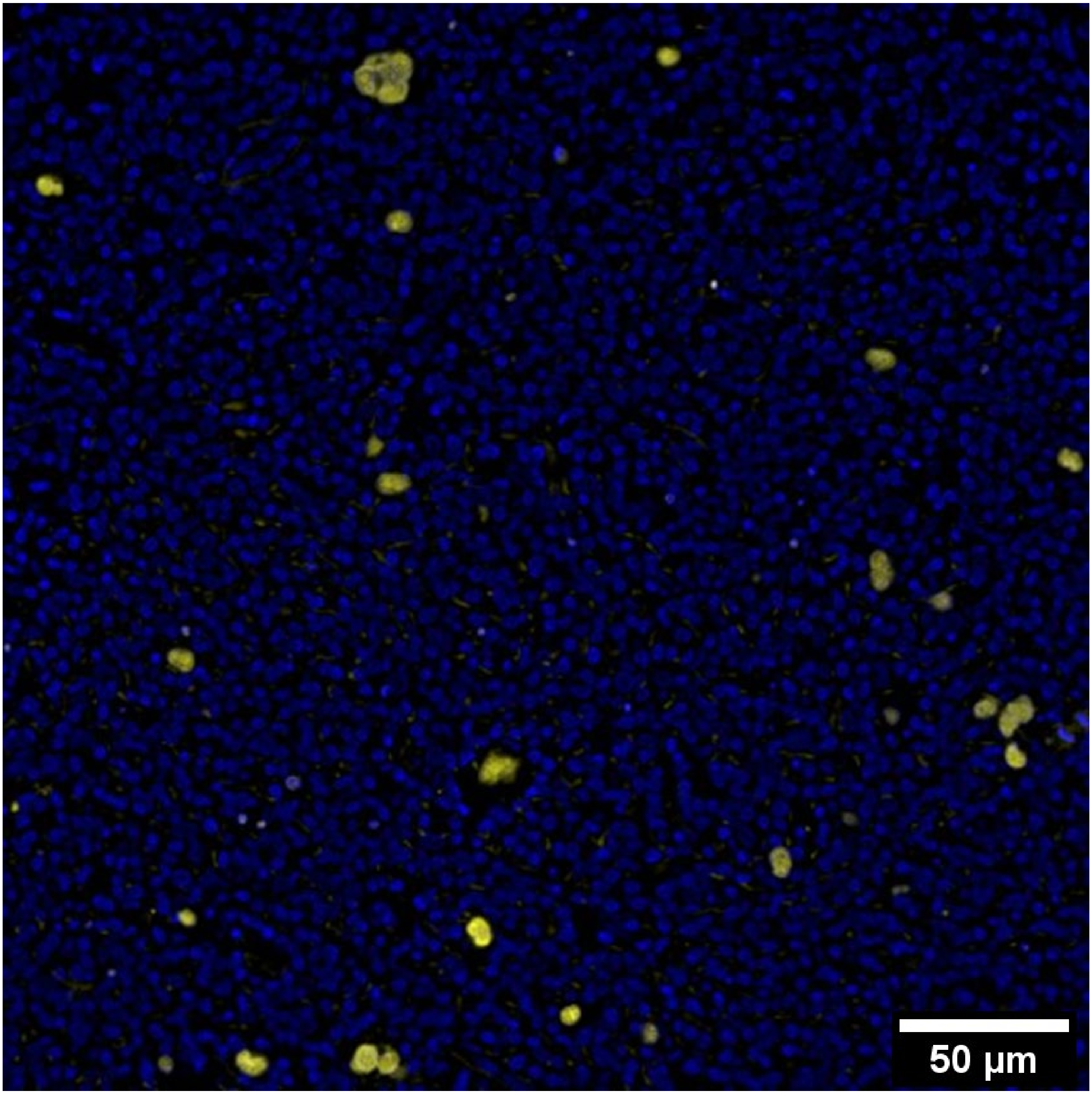


Figure S11. EBER *in situ* hybridization (ISH) imaging using tyramide-barcode staining. Staining was performed on a Leica BondRX autostainer using EBER ISH probes conjugated to FITC, followed by sequential incubation with anti-FITC primary antibodies, secondary antibodies, peroxidase conjugated tertiary antibodies, and custom tyramide-barcodes. Imaging was performed with DAPI (shown in blue) and Cy5-conjugated complementary DNA barcodes (shown in yellow). Positive EBER staining is appropriately seen in nuclei of infiltrating large cells, consistent with EBV+ Hodgkin/Reed-Sternberg cells; some weak interstitial staining is also seen. Camera/microscope settings: 500 ms exposure in Cy5 channel, 30% excitation intensity, "high sensitivity" imaging, Nikon 20x PlanApo lambda 0.75 NA objective.

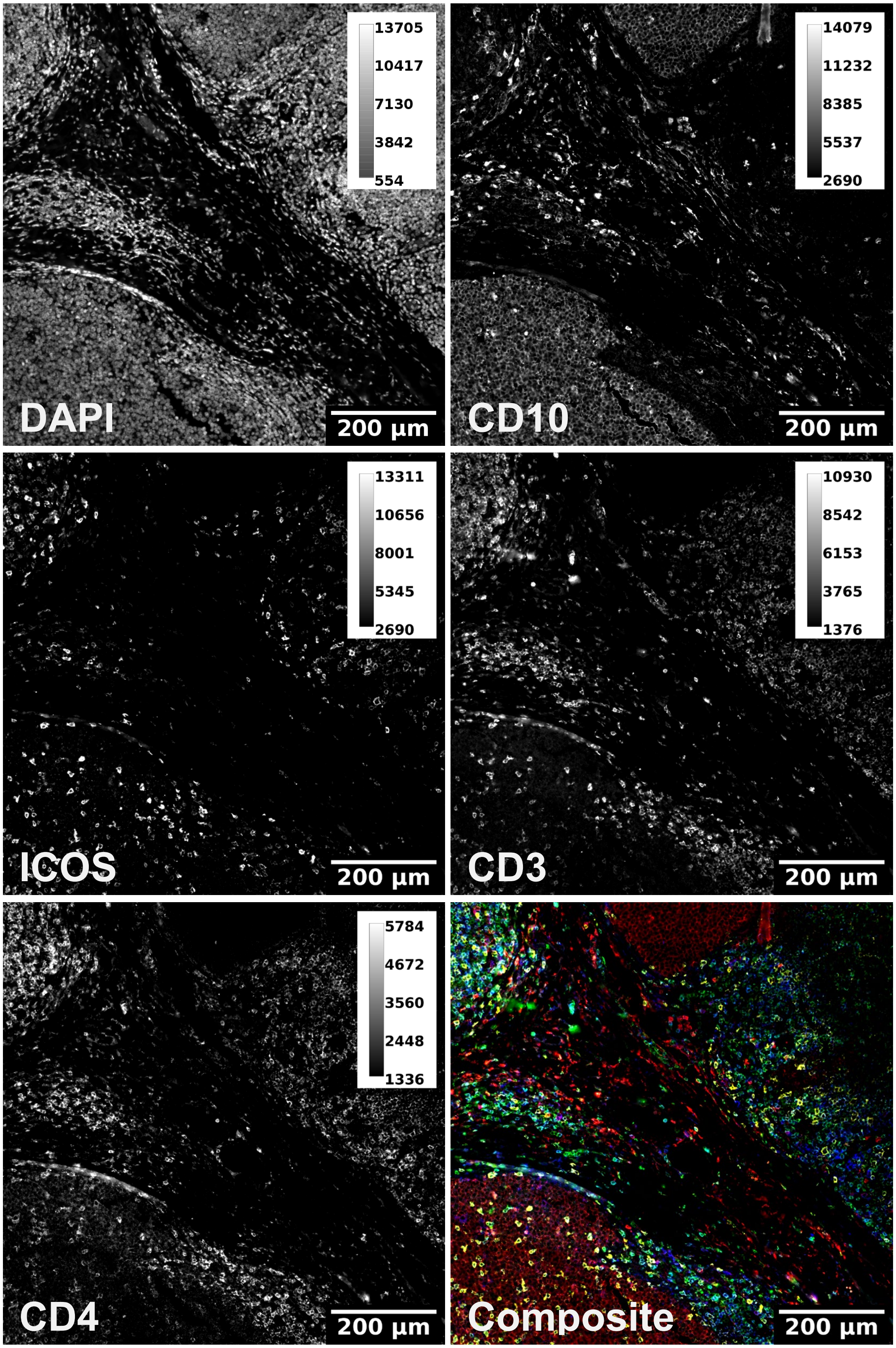


Figure S12. Multiple antibody staining using four tyramide-barcodes in tonsil, imaged using an Akoya CODEX instrument and Keyence BZ-X800 microscope. Tissue was stained using a Leica Bond RX autostainer with rabbit CD3e (1:100 dilution, clone SP7, Sigma), mouse CD4 (1:100 dilution, clone 4B12, Leica cat. No. NCL-L-CD4-368), mouse CD10 (1:200 dilution, clone OTI2A4, Origene cat no. CF810614) and rabbit ICOS (1:200 dilution, clone D1K2T, Cell Signaling cat. No.

89601) primary antibodies with signal amplification using tyramide-barcodes as described in above. The tissue was then imaged on the CODEX/Keyence system with Cy5-conjugated complementary DNA barcodes and 500 ms exposure time. The composite image is colored as follows: CD10 = red, ICOS = yellow, CD3e = green, and CD4 = blue. The Keyence BZ-X800 microscope used 100% excitation intensity, “high resolution” imaging, and a Nikon 20x PlanApo lambda 0.75 NA objective. Background subtraction and image stitching were performed using Akoya’s CODEX Processor software.

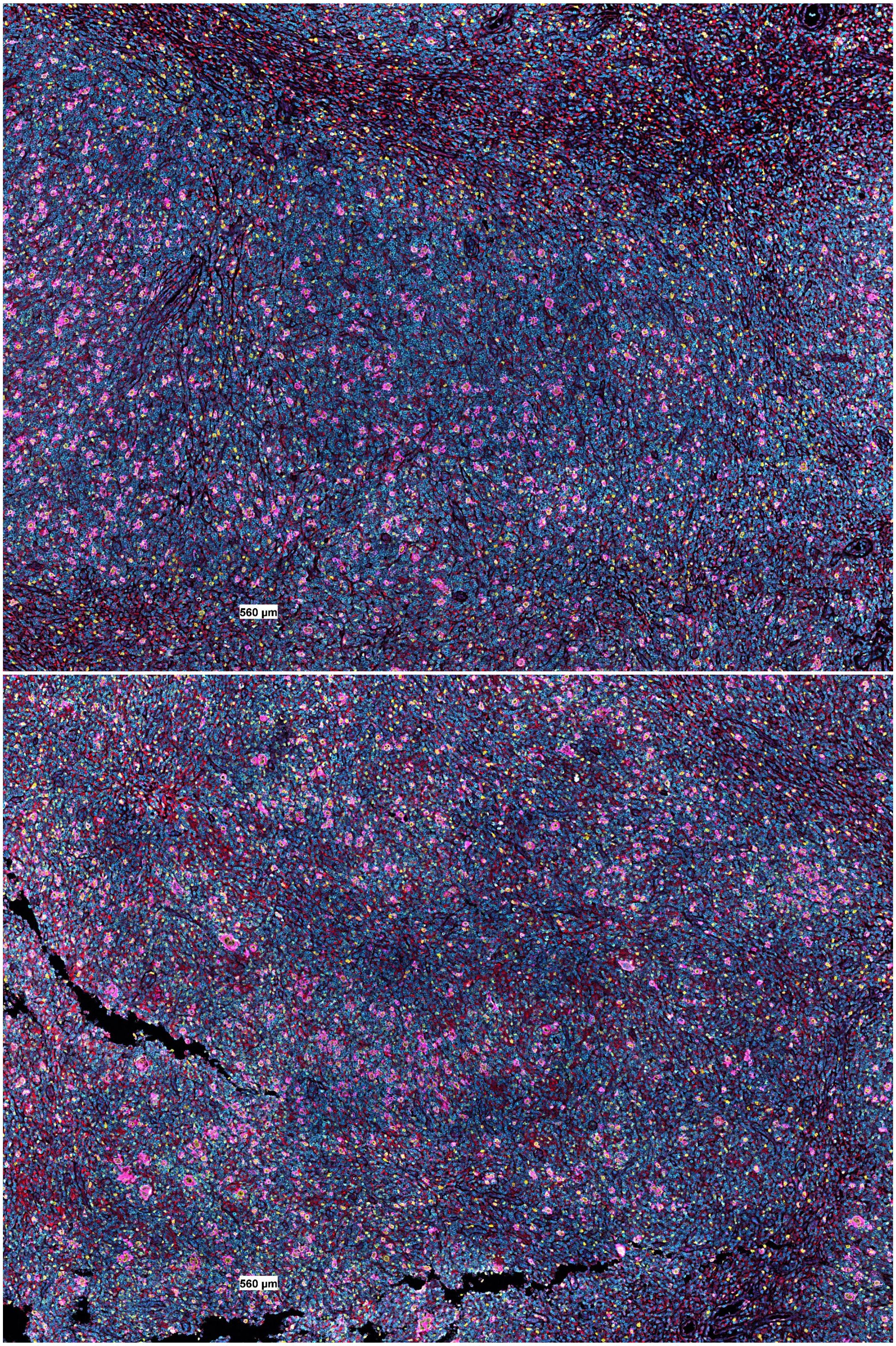


Figure S13. Overview images of classic Hodgkin lymphoma stained with commercially available barcode-conjugated antibodies and CD30 and MUM1 stained using tyramide barcodes. The top (region 1) and bottom (region 2) images are the two complete regions captured by CODEX imaging. DAPI = blue, CD30 = magenta, MUM1 = yellow, CD68 = red, CD3e = cyan. See also Fig. 3, which is composed of a zoomed-in region from the top image, and Fig. S14.

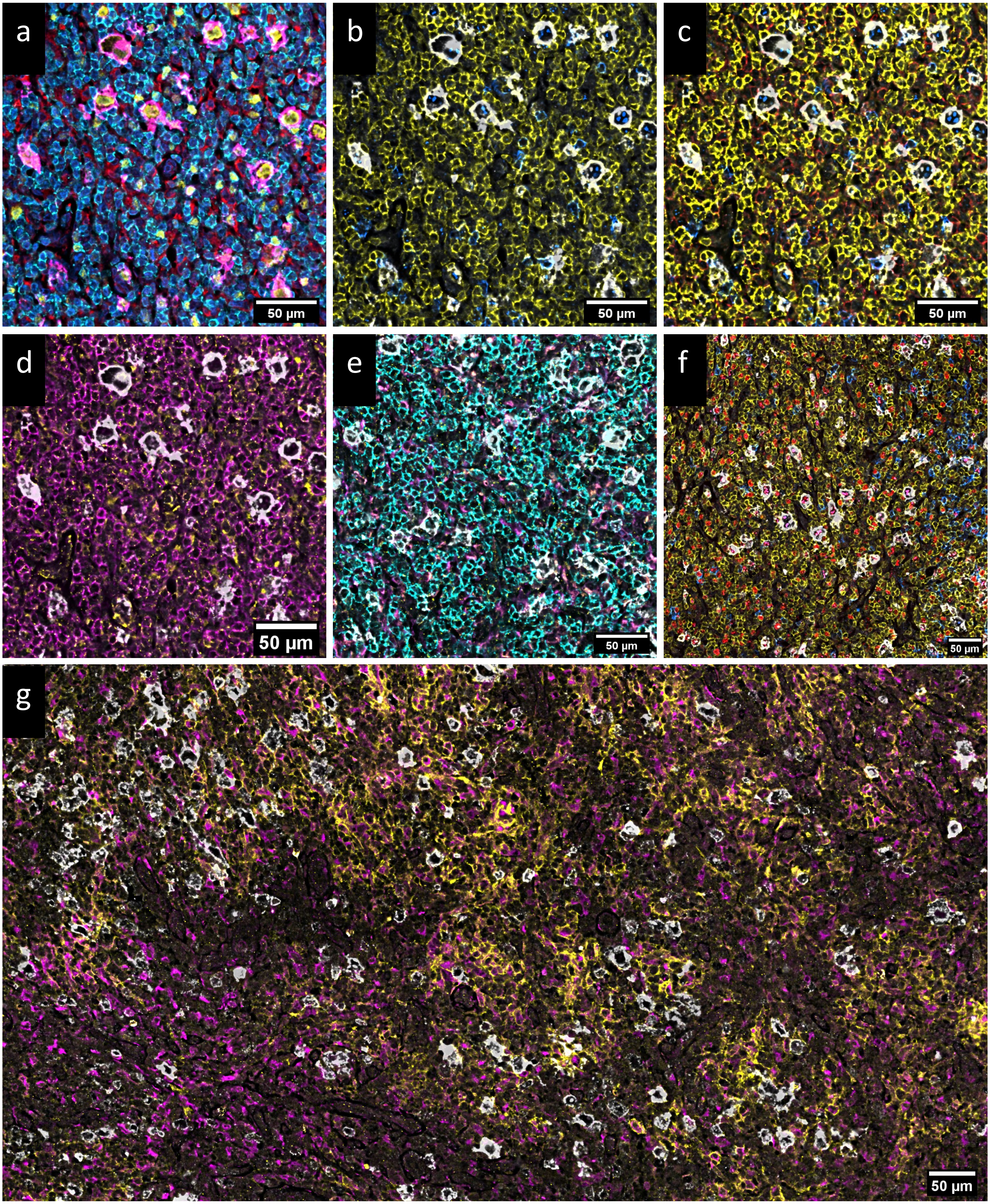
a
50 µm
b
50 µm
c
50 µm
d
50 µm
e
50 µm
f
50 µm
g
50 µm

Figure S14. Additional images of classic Hodgkin lymphoma demonstrating the various stains performed. **a:** CD30 = magenta, MUM1 = yellow, DAPI = blue, CD3e = cyan, CD68 = red. **b:** CD30 = white, CD3 = yellow, CD20 = blue. **c:** CD30 = white, CD8 = yellow, CD4 = red, CD20 = blue. **d:** CD30 = white, CD107a = yellow, CD45RO = magenta. **e:** CD30 = white, CD68 = magenta, CD107a = yellow, CD8 = cyan. CD107a is seen variably expressed in CD68+ histiocytes. CD107a puncta are also seen in association with CD8+ cells, though some other puncta are also seen which are of uncertain significance and might indicate non-specific staining. **f:** CD30 = white, CD3e = yellow, CD20 = blue, Ki67 = red. **g:** CD30 = white, CD68 = magenta, CD11c = yellow. CD11c expression is distributed in a patchy manner in the imaged tissue and primarily found in region 2 (compare with Fig. S13; a CD11c rich area is shown here) and associated more frequently with helper T cells and histiocytes. Images were created using the Akoya CODEX MAV plugin for ImageJ.

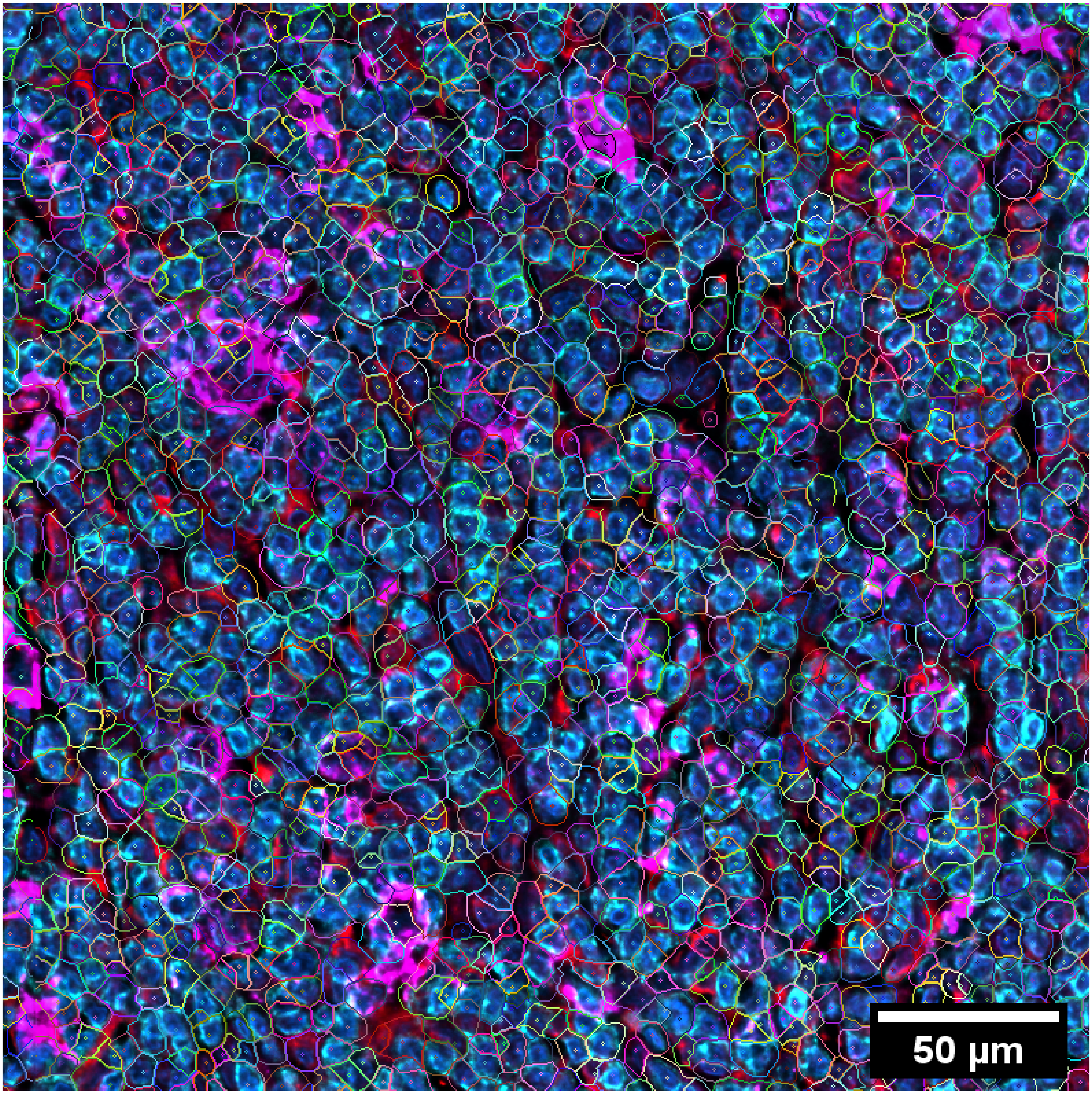


Figure S15. Example cell segmentation in case of classic Hodgkin lymphoma. Image segmentation into cells was performed using Akoya Bioscience's CODEX Processor software and displayed here using the CODEX MAV plugin (Akoya Biosciences) for ImageJ. Cell segmentation is noted to be imperfect, particularly in the large Hodgkin cells, highlighted by CD30 staining, which are subdivided into smaller Hodgkin cell segments. DAPI = blue, CD30 = magenta, CD68 = red, CD3e = cyan.

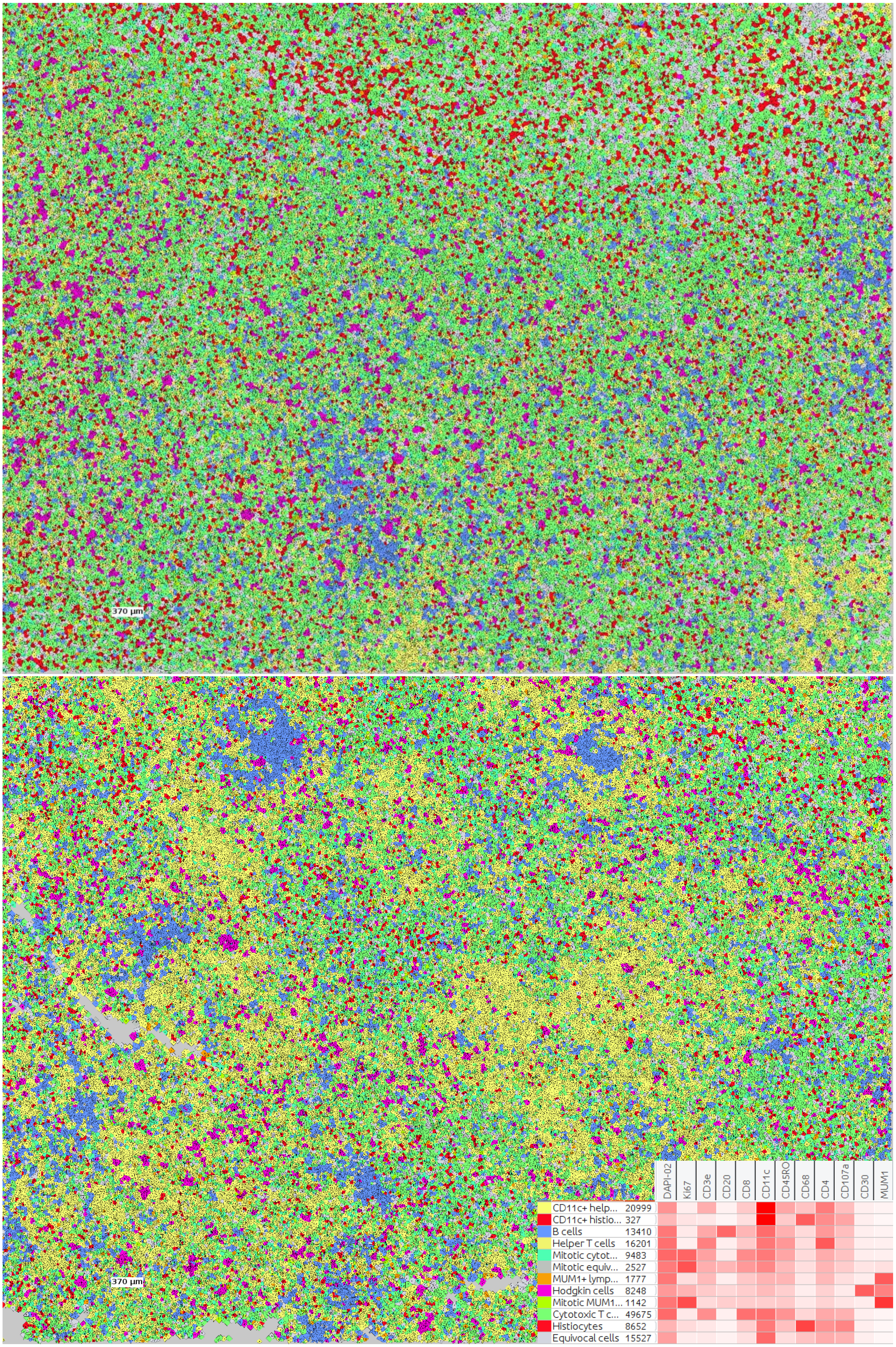
370 µm
370 µm
DAPI-02
Ki67
CD3e
CD20
CD8
CD11c
CD45RO
CD68
CD4
CD107a
CD30
MUM1
CD11c+ help... 20999
CD11c+ histio... 327
B cells 13410
Helper T cells 16201
Mitotic cytot... 9483
Mitotic equiv... 2527
MUM1+ lymp... 1777
Hodgkin cells 8248
Mitotic MUM1... 1142
Cytotoxic T c... 49675
Histiocytes 8652
Equivocal cells 15527

Figure S16. Voronoi plots with cell classifications in both imaged regions of the example case of classic Hodgkin lymphoma (see also Fig. S13). Cell classifications were determined by using the X-shift algorithm implemented in the CODEX MAV plugin (Akoya Biosciences) for ImageJ, followed by manually combining cell clusters that are of similar immunophenotype and morphologic findings. Cells labeled as “CD11c+ helper T cells” and “CD11c+ histiocytes” are in fact most likely to be helper T cells and histiocytes that do not express CD11c but are adjacent to CD11c+ dendritic cells. Cells labeled as “equivocal” appear largely to be composed, among other cell types, of endothelial cells and stromal cells. Voronoi plots are produced using the CODEX MAV plugin.

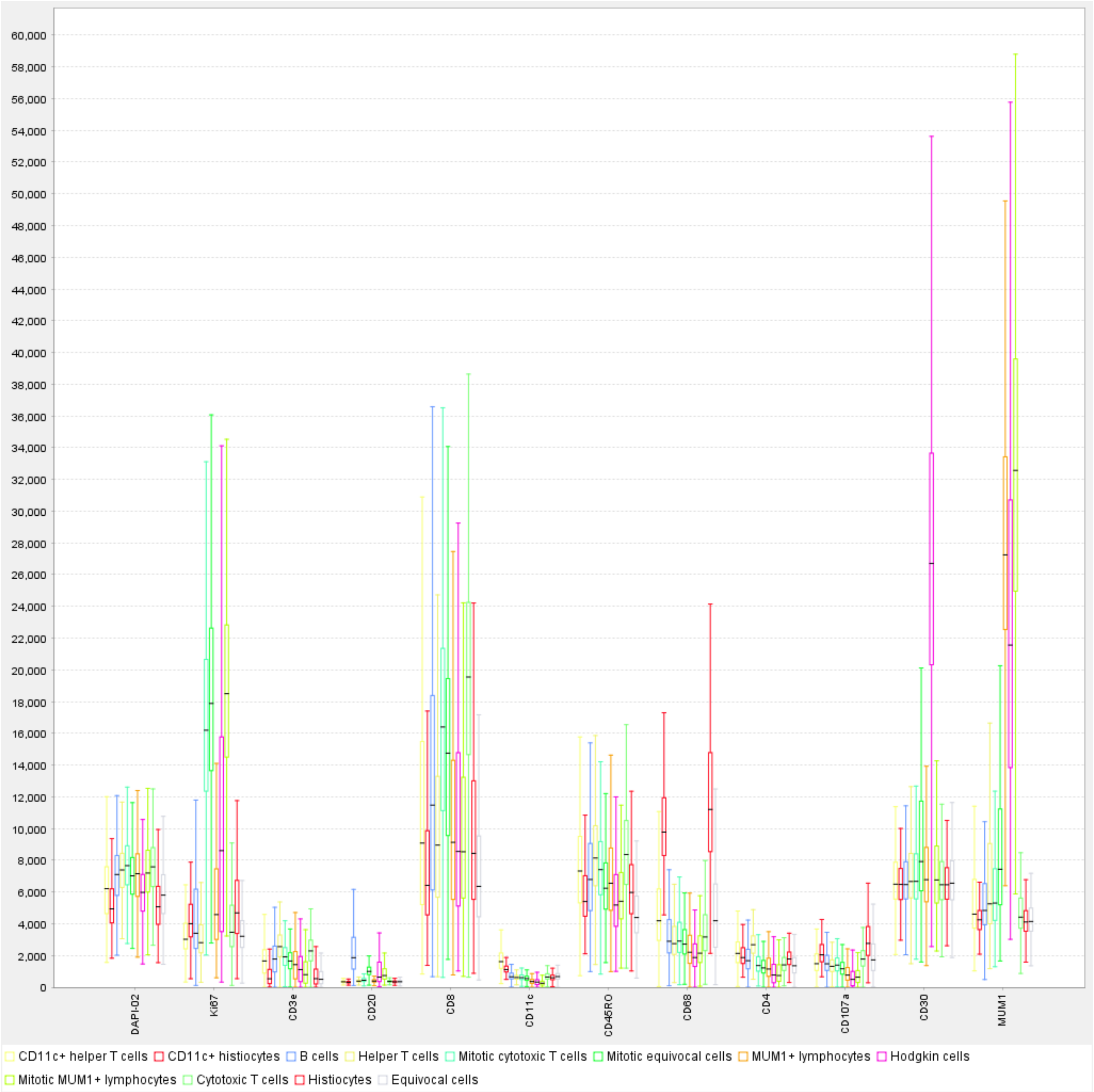


Figure S17. Box plot presentation of staining intensities for cells in tissue involved by classic Hodgkin lymphoma (see also Figs. 3, S13, and S14). Boxes indicate upper and lower quartiles; lines extending beyond the boxes (whiskers) demonstrate additional variability above and below the upper and lower quartiles. Of note are the very strong relative intensities of the CD30+ Hodgkin cells and MUM1+ cells (Hodgkin cells and some lymphocytes), attributed to tyramide signal amplification.

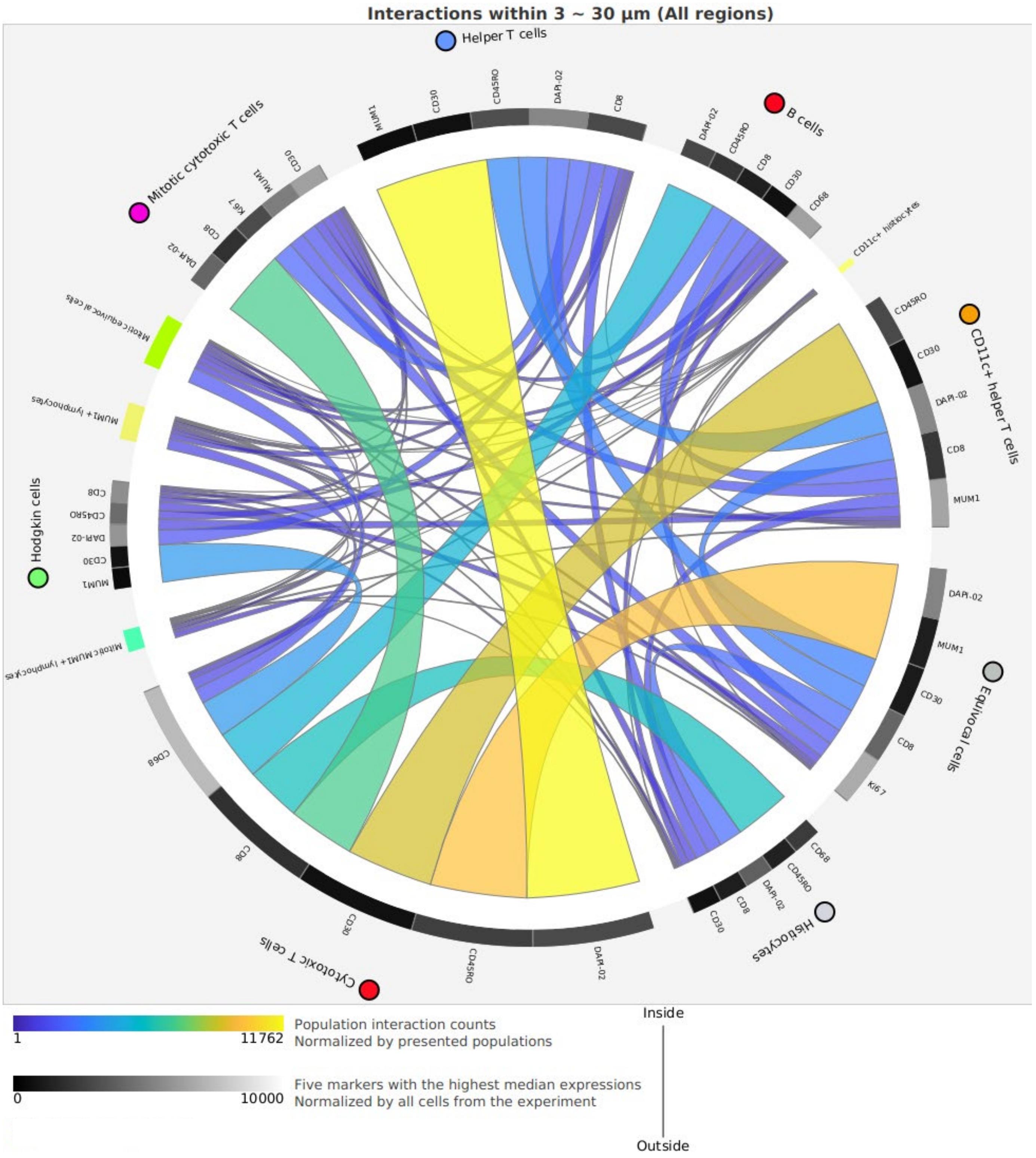


Figure S18. Circos plot of spatial analysis of nearby cells in the example case of classic Hodgkin lymphoma (both imaged regions). The Circos plot was created using the CODEX MAV plugin (Akoya Biosciences) for ImageJ. DAPI-02 refers to DAPI signal in the second CODEX imaging cycle. The five markers with highest median expressions is somewhat misleading since the scale of the intensities of the various markers differ significantly (compare with Fig. S17).
